# Multidimensional Single-Cell Transcriptomic Profiling of Uterine Leiomyosarcomas Reveals Distinct Tumor States with Inferred Therapeutic Vulnerabilities

**DOI:** 10.1101/2025.11.17.688894

**Authors:** Maria Korah, James P. Agolia, Biren Reddy, Renceh A.B. Flojo, Amanda Gonçalves, Lilin Wang, Rishik Bethi, Wesley Bobst, Brooke Liang, Kaylin A. Yip, Angela Tabora, Chia-Hsin Hsu, Rosyli Reveron-Thornton, Chuner Guo, Robert F. Gruener, Andrea E. Delitto, Peter Y. Xie, Antonio Tomasso, Beatrice Sun, Sara K. Daniel, Babak Litkouhi, Oliver Dorigo, Nam Q. Bui, Minggui Pan, Everett J. Moding, Anusha Kalbasi, Amanda R. Kirane, George A. Poultsides, Jeffrey A. Norton, Aaron M. Newman, Gregory Charville, Kristen N. Ganjoo, Deshka S. Foster, R. Stephanie Huang, Byrne Lee, Michael T. Longaker, Daniel Delitto

## Abstract

Uterine leiomyosarcoma (ULMS) is an orphan disease that frequently recurs and metastasizes, with patients undergoing multiple lines of chemotherapy due to lack of effective therapeutic targets. To address this gap, we used single-cell RNA sequencing and spatial transcriptomic analysis to comprehensively profile ULMS. We uncovered multiple states of tumor cells, including tumor cells with mesenchyme-like features, ischemic tumor cells defined by a MYC program, inflammatory tumor cells with active interferon signaling, and stem cell-like hormone receptor-positive cells. The inferred spatial correlates of these tumor cell states demonstrated unique localization patterns. By correlating these signatures to bulk RNA sequencing data, we demonstrate the relevance of these findings to clinical outcomes. Finally, using the single-cell integration and drug response prediction algorithm (scIDUC), we propose drug predictions that may target specific tumor states. Our findings suggest new avenues for further exploration of individualized and multifaceted therapeutic strategies to treat ULMS.

## Main

Uterine sarcomas account for only 3-7% of all uterine cancers^1–4^ but have disproportionate morbidity and mortality. The dominant subtype, uterine leiomyosarcoma (ULMS), comprises about 70% of all uterine sarcomas,^3^ with an annual incidence of less than one per 100,000.^5^ Even for patients diagnosed with stage I disease, the five-year overall survival is 57%.^6^ ULMS is also known for its high recurrence rates, estimated at 40-70%.^7–11^ This is in part due to diagnostic challenges: ULMS is often difficult to diagnose pre-operatively, limited by a lack of specific biomarkers or pathognomonic imaging findings to distinguish them from benign uterine leiomyomas, or fibroids. When these patients undergo minimally invasive procedures for presumed benign disease, mechanical fragmentation of the tumor for removal through small incisions can inadvertently disseminate undiagnosed uterine sarcoma throughout the abdomen, converting localized disease into widespread metastases.^12^ For patients with advanced stage disease, treatment plans are individualized but include combinations of surgery, radiation, and chemotherapy. Doxorubicin or a combination of gemcitabine and docetaxel are commonly used first-line chemotherapy agents,^6,13^ although both regimens show limited efficacy.^14^ While multiple clinical trials for advanced disease are underway, response rates remain low, and early closure due to slow accrual rates has been an issue.^15,16,17,18^

An additional hindrance to effective therapies for ULMS is our incomplete understanding of the pathophysiology driving the disease. Genomic profiling has consistently shown *TP53*, *RB1*, *ATRX*, *PTEN*, and *MED12* to be frequently mutated in ULMS.^19–22^ However, many of these mutations cannot be targeted easily, which has led researchers to bulk transcriptomic approaches. Tumor subtyping based on bulk transcriptomic studies has revealed that ULMS tends to segregate away from other leiomyosarcomas, and translation of these findings has been limited.^23–26^ For example, some tumors have a gene signature suggestive of homologous recombination deficiency;^19,27^ however, clinical trials of olaparib based on this finding were initially promising but have since stalled.^28,29^ As bulk transcriptomic studies may be limited by the heterogeneity of these tumors, single-cell resolution analyses may help to derive better targeted treatments for these patients.

In this study, we present a comprehensive single-cell transcriptomic atlas of ULMS. Using both 10x Genomics single-cell RNA sequencing (scRNA-seq) of fresh ULMS tumors collected at the time of surgical resection, and Singular G4X spatial transcriptomic analysis on paired samples, we uncover several tumor cell states with unique inferred drug sensitivities and patient outcomes. Our work refines the molecular understanding of ULMS heterogeneity and its clinical correlates at single-cell resolution, which may further inform combinatorial and personalized adjuvant therapy options.

## Results

### Patients with ULMS have complex treatment courses and genetically distinct tumors

To explore the transcriptomic landscape of ULMS, 18 patients were recruited, and fresh tumor specimens were collected at the time of surgery for recurrent disease (**Fig. 1a, b, and Supplementary Table 1, 2**). Most tumors were reported positive for estrogen receptor (ER) by immunohistochemical staining on routine pathology, and fewer tumors were positive for progesterone receptor (PR). Some tumors had changes in their hormone receptor expression pattern across successive resections in the same patient. For example, Patient 7 had decreased expression of both ER and PR on their fifth surgery compared to their second. Though the exact mechanisms driving these changes are not known, these findings may be reflective of chemotherapy-mediated clonal selection in tumor states, along with tumor plasticity and evolution. These findings may also be artifactually driven by sampling bias and a reflection of intratumoral heterogeneity of hormone receptor expression patterns, which has been extensively described in other tumors like breast and prostate cancer.^30,31^ Final pathology for all patients in our cohort was reported as leiomyosarcoma, and this included three myxoid and two epithelioid leiomyosarcoma samples. We collected two samples from five patients (Patients 1, 2, 7, 13, and 14) who had additional recurrences during the sample collection period.

**Figure 1.**
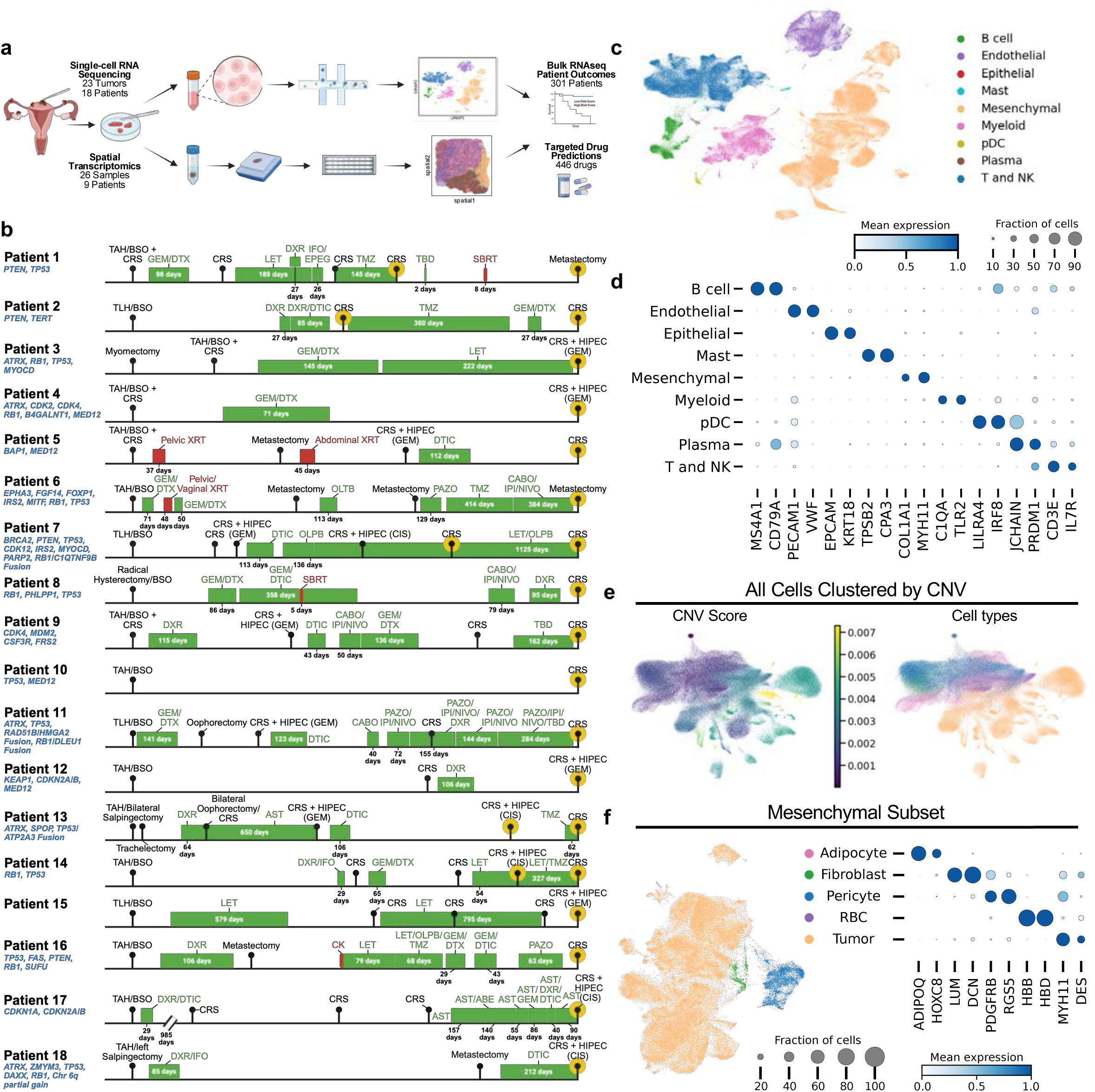
Study overview and scRNA-seq atlas of ULMS tumors. **a,** Schematic of study design integrating single-cell RNA sequencing (scRNA-seq) and spatial transcriptomic profiling. **b,** Clinical timelines of the 18 patients whose samples were analyzed in this study, showing their treatment history and surgeries prior to sample collection. Green bars represent chemotherapy, red bars indicate radiotherapy, and black circles denote surgeries, with sample collection highlighted in yellow. Mutations from clinical testing are shown in blue. Chemotherapy abbreviations are defined in **Supplementary Table 2**. TAH – total abdominal hysterectomy; BSO – bilateral salpingo-oophorectomy; CRS – cytoreductive surgery; SBRT – stereotactic body radiation therapy; TLH – total laparoscopy hysterectomy; HIPEC – hyperthermic intraperitoneal chemotherapy; XRT – radiotherapy. **c,** Batch-corrected Uniform Manifold Approximation and Projection (UMAP) of all cells following integration, colored by cell type. **d,** Dot plot showing mean expression of canonical marker genes across cell types. **e,** Batch-corrected UMAP of all cells clustered by CNV, colored by CNV score (left) or annotated cell type (right). The color scheme for the annotated cell types matches the legend in Fig. 1c. **f**, Batch-corrected UMAP of mesenchymal cells subsetted, re-integrated, clustered, and colored by cell type (left), along with the associated dot plot showing mean expression of canonical markers across cell types (right).

No two patients received the exact same therapeutic regimen, highlighting the lack of consistent and effective therapies for ULMS. Patients in this study received varying combinations, durations, and timing of chemotherapy, radiation, and cytoreduction, with each regimen tailored to individual patient factors. Most patients cycled through multiple systemic agents, including one patient who received nine different chemotherapy agents prior to sample collection. These heterogeneous treatment regimens further highlight the lack of effective therapies for ULMS.

Based on clinically available tumor genotyping results, there were no recurrently mutated genes or patterns found across all patients, including variants of unknown significance; this is consistent with prior genomic studies of ULMS (**Fig. 1b, Extended Data Fig. 1, 2, and Supplementary Table 3**).^19–22^ Even among patients with known pathogenic mutations in the same genes, the exact variants were different. The most commonly mutated genes of known significance found in our patient cohort were *TP53*, *RB1*, *ATRX*, *PTEN*, and *MED12*, consistent with prior genomic studies of ULMS.^19–22^ Several genes with variants of unknown significance were consistently observed, such as *KCNRG* and *TRIM13*, though the significance of these variants remains unclear. Altogether, these findings highlight the phenotypic and genetic heterogeneity of ULMS and the urgent need for more efficacious treatment options for this disease.

### Single-cell RNA sequencing of ULMS reveals a heterogeneous tumor microenvironment

Given the genomic heterogeneity across our ULMS tumor cohort, we sought to delineate the degree of transcriptomic heterogeneity. To accomplish this, scRNA-seq was performed on 23 tumor specimens from 18 patients. After applying standard quality control metrics, integration was performed across samples and batches using scVI, which appeared to adequately account for batch effect when pre- and post-integration UMAPs were compared (**Extended Data Fig. 3a-d**).^32^ Single-cell integration benchmarking (scIB) further revealed that scVI outperformed other methods tested, namely Harmony and no integration, in both batch correction and biological conservation (**Extended Data Fig. 3e**).^33,34^ Following pre-processing and integration, the 23 specimens yielded a total of 223,655 cells (**Fig. 1c, d**). All tumors were composed of multiple cell types, including myeloid cells, lymphoid cells, endothelial cells, and mesenchymal cells, revealing a rich tumor microenvironment. Mesenchymal cells, which included tumor cells, pericytes, and fibroblasts, were the predominant cell type in the overall cohort (102,102 cells, 45.7%).

InferCNV was then used to compute copy number variation (CNV) inferences within all cells.^35,36^ In this method, CNV score is defined as the scaled log fold change per chromosomal location relative to a reference cell type, and the mean of the absolute value of CNV scores is taken per cluster, such that a high CNV score suggests aberrant CNV in a given cluster.^35,36^ The mesenchymal cells had a higher number of CNV aberrations across chromosomes when compared to other cell types (**Extended Data Fig. 4a, 5**). Additionally, when clustered by CNV features, the mesenchymal cells clustered separately from all other cells (**Fig. 1e**). Clustering by CNV also segregated the mesenchymal cells by sample, capturing the varying clonal origins of these cells (**Extended Data Fig. 6a**).

Given the mesenchymal origins of leiomyosarcoma tumor cells, it was challenging to separate out the tumor cells from other mesenchymal cells. To facilitate this distinction, the mesenchymal cells were subsetted, re-integrated, clustered, and annotated. The tumor cells clustered separately from the fibroblasts and pericytes (**Fig. 1f**). Additionally, tumor cells had the highest CNV score of all cells, whereas cells identified as fibroblasts and pericytes based on canonical marker expression had lower CNV scores (**Extended Data Fig. 6b**). To further confirm the fibroblast and tumor cell distinction, a composite score encompassing markers previously shown to delineate fibroblasts from soft tissue sarcoma tumor cells was plotted, which further validated these annotations (**Extended Data Fig. 6c**). Altogether, these findings highlight the heterogeneous tumor microenvironment of ULMS.

### ULMS tumor cells demonstrate heterogenous transcriptomic states with three prevalent mechanisms

Given the known heterogeneity of ULMS tumors, we were interested in defining single-cell tumor states. After reintegrating and re-clustering the tumor cells, we identified nine clusters (**Fig. 2a, b**). These clusters were largely stable across Leiden and Louvain clustering algorithms, but since the Leiden clusters appeared more biologically meaningful based on top differentially expressed genes, the remainder of the analyses were based on the Leiden clusters (**Extended Data Fig. 7a, b**). These tumor clusters were distributed in varying proportions across all samples, with no single patient sample solely contributing to any one tumor cell state (**Fig. 2c**).

**Figure 2.**
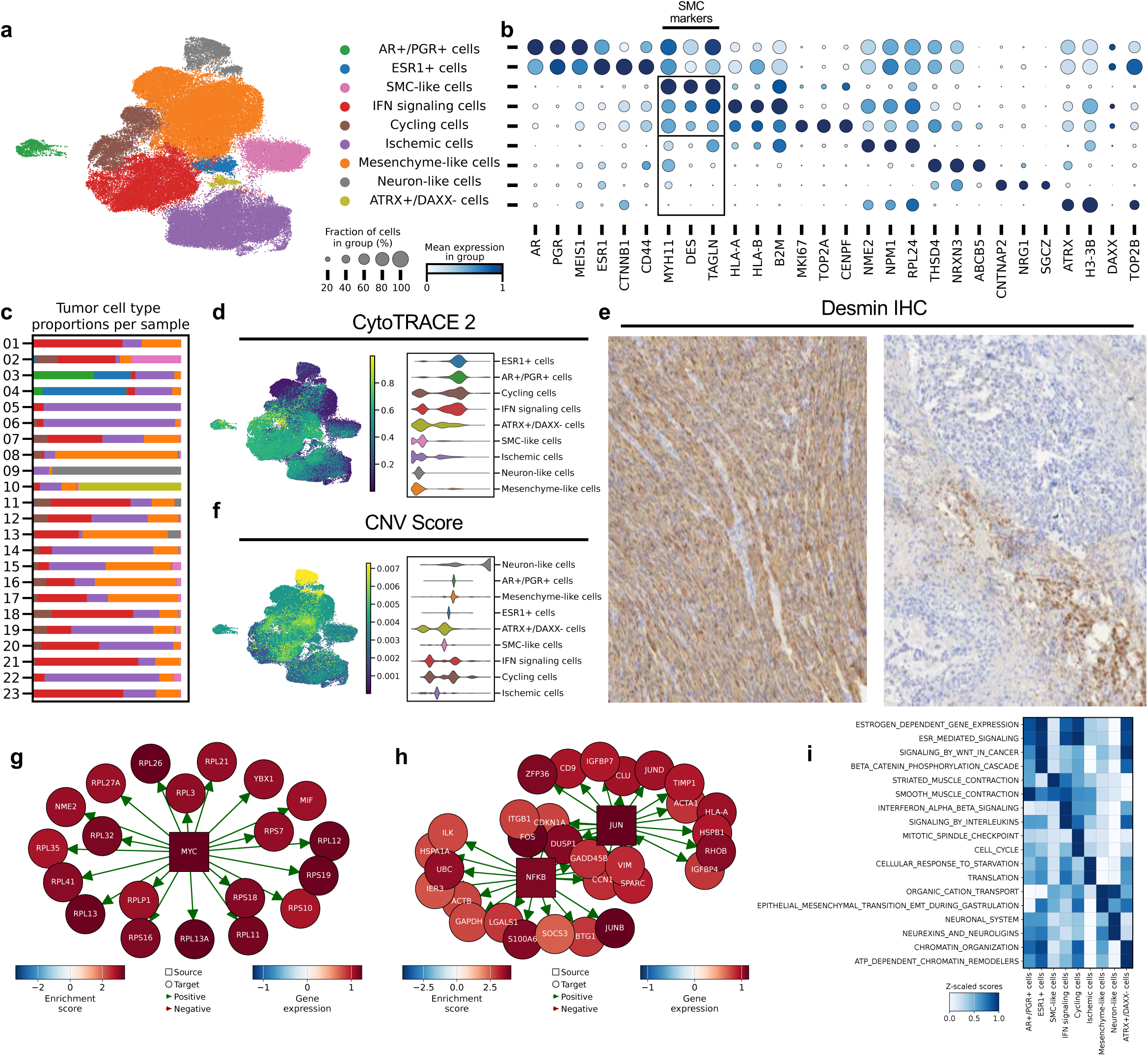
ULMS tumor cell states delineated with scRNA-seq. **a,** Batch-corrected UMAP of subsetted, re-integrated, and clustered tumor cells annotated by tumor cell state. **b,** Dot plot showing mean expression of selected markers for each tumor cell state. **c,** Proportion of tumor cell states in each patient sample. **d,** Batch-corrected UMAP colored by CytoTRACE 2 cellular potency scores, indicating inferred differentiation potential (yellow = higher potency; blue = lower potency). Quantification of pre-KNN CytoTRACE 2 scores for each tumor state (right). **e,** Representative immunohistochemistry (IHC) images for desmin (DES), one with high (left) expression of DES and the other with relatively lower (right) expression of DES. **f,** Batch-corrected UMAP colored by InferCNV score (left) (yellow = higher CNV score; blue = lower CNV score) with corresponding quantification (right). **g,** Network diagram highlighting top genes from representative regulatory modules for MYC identified from CollecTRI regulon inference for Ischemic cells. **h,** Network diagram highlighting top genes from representative regulatory modules for JUN and NFKB identified from CollecTRI regulon inference for IFN signaling cells. **i,** Heatmap of selected Reactome pathways enriched in each tumor state.

Two of the clusters had high expression of markers implying hormonal signaling, including androgen receptor (*AR*), progesterone receptor (*PGR*), and estrogen receptor (*ESR1*) (**Fig. 2a, b, Extended Data Fig. 7c, and Supplementary Table 4**). Interestingly, these cells separated into two clusters: (1) those high in *AR* and *PGR* gene expression but lower in *ESR1* expression (termed AR+/PGR+ cells) and (2) those that have high *ESR1* but lower *AR* and *PGR* (termed ESR1+ cells). The ESR1+ cells were also enriched in markers of Wnt pathway signaling (such as *CTNNB1*) and myometrial stem cell markers (such as *CD44*).^37–40^ In contrast, the AR+/PGR+ cells had lower enrichment of Wnt pathway signaling markers and *CD44* but had high expression of other developmental pathway markers such as *MEIS1*.^41,42^ Additionally, the ESR1+ cells had relatively lower expression of canonical smooth muscle cell (SMC) markers (such as *MYH11*, *DES*, and *TAGLN*), whereas the AR+/PGR+ cells had relatively higher expression of some of these canonical SMC markers (**Extended Data Fig. 7d**).

The gene expression signatures of the ESR1+ cells and AR+/PGR+ cells in ULMS tumors were suggestive of a less differentiated, stem cell-like phenotype. In fact, myometrial stem cells have previously been described as having similar features, with active Wnt pathway signaling, high *CD44* expression, and lower expression of SMC markers. Myometrial stem cells are also thought to be supported through paracrine hormonal signaling from surrounding mature niche cells that have higher expression of estrogen and progesterone receptors and SMC markers.^37–40^ We therefore hypothesized that the hormone receptor-expressing cells captured in our scRNA-seq dataset were more primitive, less differentiated, stem-like cells. To test this hypothesis, we used CytoTRACE 2, a deep learning model that can infer the developmental potential of each tumor cell.^43^ This analysis revealed that the ESR1+ cells and AR+/PGR+ cells were some of the least differentiated cells in the tumor subset (**Fig. 2d**). To ensure that the CytoTRACE 2 predictions were not driven by proliferation markers, we computed cell cycle scores for each cell. This analysis revealed that the ESR1+ and AR+/PGR+ cells were primarily in G1 phase, suggesting that proliferation markers were not the sole drivers of the CytoTRACE 2 predictions (**Extended Data Fig. 8a-c**). Taken together, these findings suggest that these hormonally active cells are stem cell-like.

Among the remaining non-hormonal tumor clusters, there was a distinction in the expression of SMC markers: three clusters (SMC-like cells, Interferon (IFN) signaling cells, and Cycling cells) had strong expression of canonical smooth muscle markers, such as *MYH11*, *DES*, *TAGLN*, *ACTA2*, *CALD1*, *LMOD1*, *CNN1*, and *MYLK*, while the other three clusters (Ischemic cells, Mesenchyme-like cells, and Neuron-like cells) had lower smooth muscle marker expression (**Fig. 2b, Extended Data Fig. 7d, and Supplementary Table 4**). Immunohistochemistry (IHC) was performed for desmin (DES), further confirming this variation in SMC marker expression (**Fig 2e**). To ensure that these cells expressing lower levels of canonical SMC markers were not a result of contamination with non-tumor cells, inferCNV scores were plotted for these subpopulations. Cells with decreased smooth muscle marker expression had high CNV scores, suggesting that these were tumor cells (**Fig. 2f and Extended Data Fig. 9, 10**). Higher CNV in low SMC-expressing cells suggested a potential difference in differentiation states across cells, and we hypothesized that the cells with high CNV are likely less differentiated. Conversely, CytoTRACE 2 suggested that the cells with high CNV and low SMC marker expression were not predicted to be less differentiated (**Fig. 2d**). In fact, the two clusters predicted to be the most differentiated (Mesenchyme-like cells and Neuron-like cells) also had some of the highest inferred CNV scores. Moreover, both of these clusters highly expressed markers of neuronal differentiation (*NRXN3*, *CNTNAP2*, and *NRG1*), especially the Neuron-like cells (**Fig. 2a, b, Extended Data Fig. 7c, and Supplementary Table 4**). Prior studies have identified such neuronal differentiation among rare uterine sarcomas and in the context of other sarcomas, such as Ewing’s sarcoma.^44–47^

Though both the Neuron-like cells and the Mesenchyme-like cells had high expression of markers of neuronal differentiation, the mesenchyme-like cells were distinguished by their high expression of extracellular matrix (ECM) remodeling markers, such as *THSD4* and *LARGE1*. Their fibroblast-like profile was further highlighted using SCimilarity, which queries an atlas of single cell states to generate predictions on what an input set of cells most closely resemble.^48^ SCimilarity suggested that the Mesenchyme-like cell profile is most similar to fibroblasts and myofibroblasts (**Extended Data Fig. 11a, b**). These cells also highly express *ABCB5*, an ATP-binding cassette family gene encoding a drug-efflux transporter previously shown to confer chemoresistance in other cancers (**Fig. 2a, b, Extended Data Fig. 7c, and Supplementary Table 4**).^49,50^ The high neuronal and ECM-remodeling signatures in these Mesenchyme-like tumor cells suggest that these cells may be highly plastic and associated with enhanced tumor cell migration and invasion capabilities.^51–53^ This phenotype, along with the potential for chemoresistance, may contribute to the highly invasive biology of these ULMS tumors.

The Mesenchyme-like cells appeared to be ubiquitous across almost all samples, along with two other tumor states: the Ischemic cells and the IFN signaling cells (**Fig. 2a, c**). These three tumor states were also present across samples diagnosed as myxoid (Samples 01, 09, and 11) and epithelioid (Samples 07 and 18) subtypes of leiomyosarcomas (**Extended Data Fig. 12a-d**). The presence of these three tumor cell states across samples and subtypes alluded to biological mechanisms shared across most uterine leiomyosarcomas; therefore, we sought to explore the phenotypes of these tumor subsets further.

Like the Neuron-like cells and Mesenchyme-like cells, the Ischemic cells had lower expression of canonical SMC markers (**Fig. 2a, b, Extended Data Fig. 7c-d, and Supplementary Table 4**). These cells had high expression of many ribosomal genes, such as *RPL24*. We ensured that this ribosomal biogenesis pattern reflected a biologically relevant signal and not technical artifact or low quality by verifying that these cells had strong quality control metrics, including number of genes detected in each cell, total counts from each cell, total number of mitochondrial genes in each cell, and total number of ribosomal genes in each cell (**Extended Data Fig. 13a-m**). When compared to the other tumor cell states, Ischemic tumor cells did not have the lowest genes per cell or total counts, and they did not have the highest mitochondrial or ribosomal gene counts. Additionally, the proportion of Ischemic tumor cells stayed relatively stable across samples from different time points from the same patient, further suggesting that the signature in these cells represent biologically meaningful programs (**Extended Data Fig. 13n**). Ribosomal biogenesis has previously been described to be orchestrated by MYC-mediated signaling, and these cells had high expression of *NME2* and *NPM1*, markers associated with MYC signaling pathways. To further substantiate the presence of MYC-mediated signaling in these cells, we inferred active transcription factors in all our tumor states by interrogating our dataset using the CollecTRI-derived regulon resource using the Decoupler package (**Extended Data Fig. 14a and Supplementary Table 5**).^54,55^ MYC was inferred to be highly enriched and active in these cells (**Fig. 2g and Supplementary Table 5**). MYC has been previously identified as being histologically overexpressed in a substantial number of uterine sarcomas, and its presence has been reported to be associated with worse prognoses in soft tissue leiomyosarcomas.^56,57^ MYC signaling has previously been shown to activate adaptive programs in areas of ischemia and high metabolic demand, both in the context of tumor growth and other ischemic processes like stroke and myocardial infarction, which led us to term these cells “Ischemic tumor cells.”^58–61^ MYC signaling in tumor cells is well known to promote tumor cell proliferation and growth, along with enhancing ribosomal biogenesis; its presence in ULMS suggests that high translational activity is one survival mechanism employed by these tumors.^62,63^

The third predominant tumor cell type present across most samples, the IFN signaling cells, had higher expression of canonical SMC markers compared to the Mesenchyme-like cells and Ischemic cells (**Fig. 2a, b, Extended Data Fig. 7c-d, and Supplementary Table 4**). The dominant feature of these cells was their high expression of inflammatory markers suggestive of interferon signaling (such as HLA genes and *B2M*), along with an AP-1 mediated program (such as *JUN* and *FOS*). CollecTRI-derived transcription factor inference further supported this signature: these cells were predicted to have high activity of transcription factors JUN and AP1, along with key regulators of inflammatory signaling, NFKB and RELA (**Fig. 2h and Supplementary Table 5**). These cells also have elevated expression of *CD47*, which encodes an immune checkpoint protein that when bound to SIRP-α on macrophages, facilitates evasion of phagocytosis. CD47 expression can be induced by several factors including hypoxia, interferons, and inflammatory cytokines like TNF-α and IL-1β.^64–67^ Consistent with this, these cells have higher expression of interferon gamma receptor marker *IFNGR1* and TNF signaling marker *TNFRSF1A,* along with *GADD45B*, an NFKB target gene that can modulate JNK activity levels **(Supplementary Table 5)**. TNF-α stimulation can activate parallel MAPK-regulated arms activating both NFKB and AP-1 signaling. JNK is one of several upstream drivers of the AP-1 pathway, and with modulation from GADD45B, this program can calibrate JNK activity levels to promote AP-1-mediated invasion and proliferation without apoptosis, a sequence suggestive of chronic TNF-α stimulation.^68,69^ Gene-set enrichment analysis further highlights this phenotype, with highly enriched Reactome pathways including IFN and interleukin signaling (**Fig 2i and Supplementary Table 6**).^70^ Altogether, the transcriptomic profile of these cells is suggestive of a chronic IFN and TNF-α stimulated, immune-evasive, and highly resilient tumor cell cluster.

In addition to these three ubiquitous states, some less prevalent tumor cell states were also present, with distinct biological characteristics. The Cycling tumor cells had high expression of proliferative markers such as *MKI67*, *TOP2A*, and *CENPF* (**Fig. 2a, b and Supplementary Table 4**). These Cycling cells also had transcriptomic profiles overlapping both the Mesenchyme-like tumor cells and IFN signaling cells, suggesting that these Cycling tumor cells may be the proliferating counterparts to those two prevalent clusters. Other less prevalent clusters include the Neuron-like cells, ATRX+/DAXX-that had high expression of ATRX and H3-3B, but low DAXX, suggestive of tumor cells with functional genomic instability, and SMC-like cells that had the highest expression of canonical SMC markers (**Fig. 2a, b, Extended Data Fig. 7c-d, and Supplementary Table 4**).

Overall, we identified several tumor cell states: AR+/PGR+ cells (high *AR*, *PGR*, *MEIS1*), ESR1+ cells (high *ESR1*, *CTNNB1*, *CD44*), SMC-like cells (high canonical SMC markers, such as *MYH11*, *DES*, *TAGLN*), IFN signaling cells (high HLA genes, *B2M*, *JUN*, *FOS*), Cycling cells (high *MKI67*, *TOP2A*, *CENPF*), Ischemic cells (high ribosomal genes, *NME2*, *NPM1*), Mesenchyme-like cells (high *THSD4*, *NRXN3*, *ABCB5*), Neuron-like cells (high *CTNNAP2*, *NRG1*, *SGCZ*), and ATRX/DAXX-cells (high *ATRX*, *H3-3B*; low *DAXX*). By correlating cluster-specific signatures, these tumor states were also identified in varying levels across samples from a previously published 3SEQ dataset of ULMS, suggesting that the tumor states identified using scRNA-seq here are not specific to the cohort or platform used in this study and are rather more widely applicable (**Extended Data Fig. 14a, b**). Our analyses suggest that the AR+/PGR+ cells and the ESR1+ cells represent progenitor phenotypes that could play a role in promoting and maintaining the ULMS tumor microenvironment. We also identify three predominant tumor cell states present across most samples (IFN signaling cells, Ischemic cells, and Mesenchyme-like cells) that are likely representative of three major mechanisms utilized by these highly metastatic and recurrent uterine leiomyosarcomas. The tumor states identified in this study are largely consistent with several prior bulk transcriptomic and genomic studies of leiomyosarcomas, both in expression of previously described markers and in overall inferred functional states (**Extended Data Fig. 15a-d**).

### ULMS tumors exhibit an immunosuppressive microenvironment dominated by tumor-tumor cell communication

Given that some tumor cell states were highly suggestive of an immunomodulatory influence, we sought to further understand the immune microenvironment of ULMS tumors. Myeloid cells were separately subsetted, re-integrated, re-clustered, and annotated in finer detail **(Fig. 3a, b, and Extended Data Fig. 16a)**. Among the myeloid cells, there were several cell types identified including macrophages, osteoclast-like giant cells (OGCs), neutrophils, mast cells, conventional dendritic cells (cDCs), and plasmacytoid dendritic cells (pDCs) (**Fig. 3a, b**). OGCs have previously been described in ULMS, and patients with ULMS with OGCs may have more aggressive clinical courses even with early-stage disease.^71,72^ These OGCs have high expression of *CTSK*, which encodes a protease that facilitates ECM remodeling through collagen degradation, along with *SIRPA*, which participates in the CD47-SIRPα ligand-receptor interaction that facilitates OGC formation through macrophage fusion.

**Figure 3.**
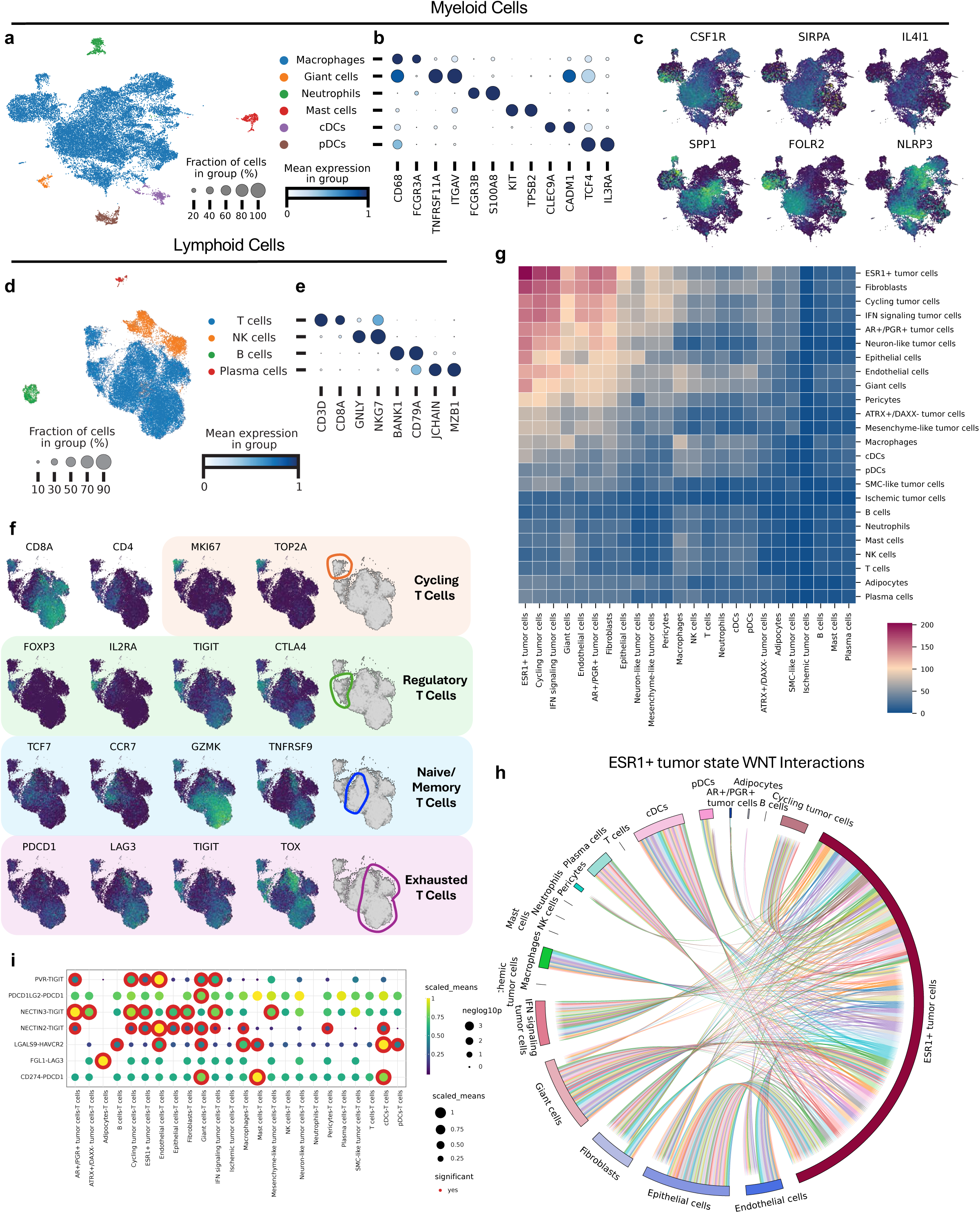
Immune cell states and intercellular communication patterns. **a,** Batch-corrected UMAP of all subsetted, re-integrated, and clustered myeloid cells, colored by cell type. **b,** Dot plot showing mean expression of selected canonical markers across annotated myeloid cell types. **c,** Batch-corrected UMAPs of macrophages colored by selected markers previously described in tumor-associated macrophages. **d,** Batch-corrected UMAP of all subsetted, re-integrated, and re-clustered lymphoid cells, colored by cell type. **e,** Dot plot showing mean expression of selected canonical markers across annotated lymphoid cell types. **f,** Batch-corrected UMAPs of T cells colored by selected markers demonstrating various T cell states. **g,** Heatmap displaying significant directional cell-cell interactions computed using CellphoneDB. **h,** CellphoneDB chord diagram displaying all Wnt-related interactions between ESR1+ tumor cells and all other cells. **i,** CellphoneDB dot plot showing selected T cell interactions.

Macrophages also displayed high expression of SIRPα (**Fig. 3c**). Stimulation of SIRPα expressed on macrophages by CD47 leads to inhibition of phagocytosis, a mechanism that was earlier highlighted in the IFN signaling tumor cells that had high expression of CD47.^67^ Macrophages also showed high expression of *CSF1R*, and stimulation of CSF1R by CSF1 secreted by tumor cells has previously been shown to promote a pro-tumorigenic phenotype in macrophages.^73,74^ These macrophages also had heterogenous expression of markers recently described in spatially distinct macrophage populations, including interleukin-4-induced gene 1 (IL4I1), secreted phosphoprotein 1 (SPP1), folate receptor beta (FOLR2), and NLR family pyrin domain containing protein 3 (NLRP3).^75^ IL4I1 macrophages have been associated with a better prognosis than SPP1 macrophages in colorectal cancer, and interestingly, in this study, macrophages had higher expression of SPP1 than IL4I1. Of note, SPP1 macrophages have previously been described to localize in areas of ischemia and induce exhaustion in CD8 T cells. While FOLR2 macrophages have been described to promote both pro-and anti-tumorigenic phenotypes, NLRP3 macrophages have been shown to promote an immunosuppressive tumor microenvironment and tumor progression.

Given that the macrophage-mediated pro-tumorigenic microenvironment maybe partly mediated by inducing T cell exhaustion, we turned to further dissecting the lymphoid microenvironment in ULMS. Lymphoid cells were separately subsetted, re-integrated, re-clustered, and annotated in finer detail **(Fig. 3d, e, and Extended Data Fig. 16b)**. Among lymphoid cells were B cells, plasma cells, NK cells, and most predominantly, T cells (**Fig. 3d, e**). Like macrophages, the T cells had heterogenous expression of markers suggestive of various functional states (**Fig. 3f**): some T cells showed high expression of markers of proliferation including *MKI67* and *TOP2A*, suggestive of cycling and actively proliferating T cells. Another population of T cells had high expression of markers associated with regulatory T cells, including *CD4*, *FOXP3*, *IL2RA*, *TIGIT*, and *CTLA4*. There were also T cells with naïve and memory T cell markers such as *TCF7* and *CCR7*. Some of these naïve and memory T cells had high expression of *GZMK* and *TNFRSF9*, while others did not, suggestive of a mixed population of both naïve and memory T cells. Finally, a large group of T cells had high expression of *CD8A* and markers of exhaustion, including *LAG3*, *TIGIT*, *PDCD1*, and *TOX*, further highlighting an immunosuppressed ULMS microenvironment. Overall, these findings suggest that both myeloid and lymphoid cells in ULMS coordinate and promote an immunosuppressed and pro-tumorigenic microenvironment. This is consistent with prior studies of the ULMS immune microenvironment, including a recent study looking at scRNA-seq of ULMS tumors across time from one patient, where they also identified a predominantly immunosuppressed tumor microenvironment.^76^

Given the heterogeneity in cell states amongst tumor, myeloid, and lymphoid cells, we then aimed to elucidate the intercellular communications patterns in ULMS. To do this, we used the CellphoneDB package, which predicts receptor-ligand interactions in scRNA-seq data using a publicly available database of manually curated interaction pairs.^77^ The predominant communication patterns identified were among tumor cells, between various tumor cell states, particularly ESR1+ tumor cells, AR+/PGR+ tumor cells, cycling tumor cells, and IFN signaling tumor cells (**Fig. 3g and Extended Data Fig. 16c**). ESR1+ tumor cells had the highest number of significant interactions, predominantly related to Wnt signaling and ECM remodeling, consistent with their transcriptomic phenotype described earlier (**Fig. 3g, h, and Extended Data Fig. 16a, d**).

Among immune cells, giant cells and macrophages had the most significant interactions identified, which included immunosuppressive interactions mediated by SPP1, CSF1R and SIRPα, consistent with their transcriptomic profiles (**Extended Data Fig. 16e**). The significant interactions identified among T cells also included several immunosuppressive receptor-ligand pairs, including those mediated through TIGIT, PDCD1, HAVCR2, LAG3, and PDCD1 (**Fig. 3i**). Together, these data demonstrate that ULMS tumors have a microenvironment dominated by interactions between various tumor cell states, along with an immune landscape that is primarily immunosuppressive.

### Spatial profiling of recurrent ULMS tumors uncovers varying tumor morphologies

Given the variety of mechanisms employed by the tumor states to promote invasion, proliferation, and chemoresistance, we hypothesized that the spatial localization of the tumor states would complement their transcriptional profiles. To test this hypothesis, we performed a probe-based spatial transcriptomic analysis on 26 regions from 9 of the patients in the single-cell cohort using the Singular G4X platform, which identifies expression of a panel of 309 genes with subcellular resolution.^78^ After cell segmentation with Proseg^79^ and quality control, 1,983,386 cells remained. Integration and clustering revealed several cell types, including lymphocytes, macrophages, fibroblasts, endothelial cells, pericytes, and tumor cells (**Fig. 4a, b**). Tumor cells were the predominant cell type in all sections (**Fig. 4c**).

**Figure 4.**
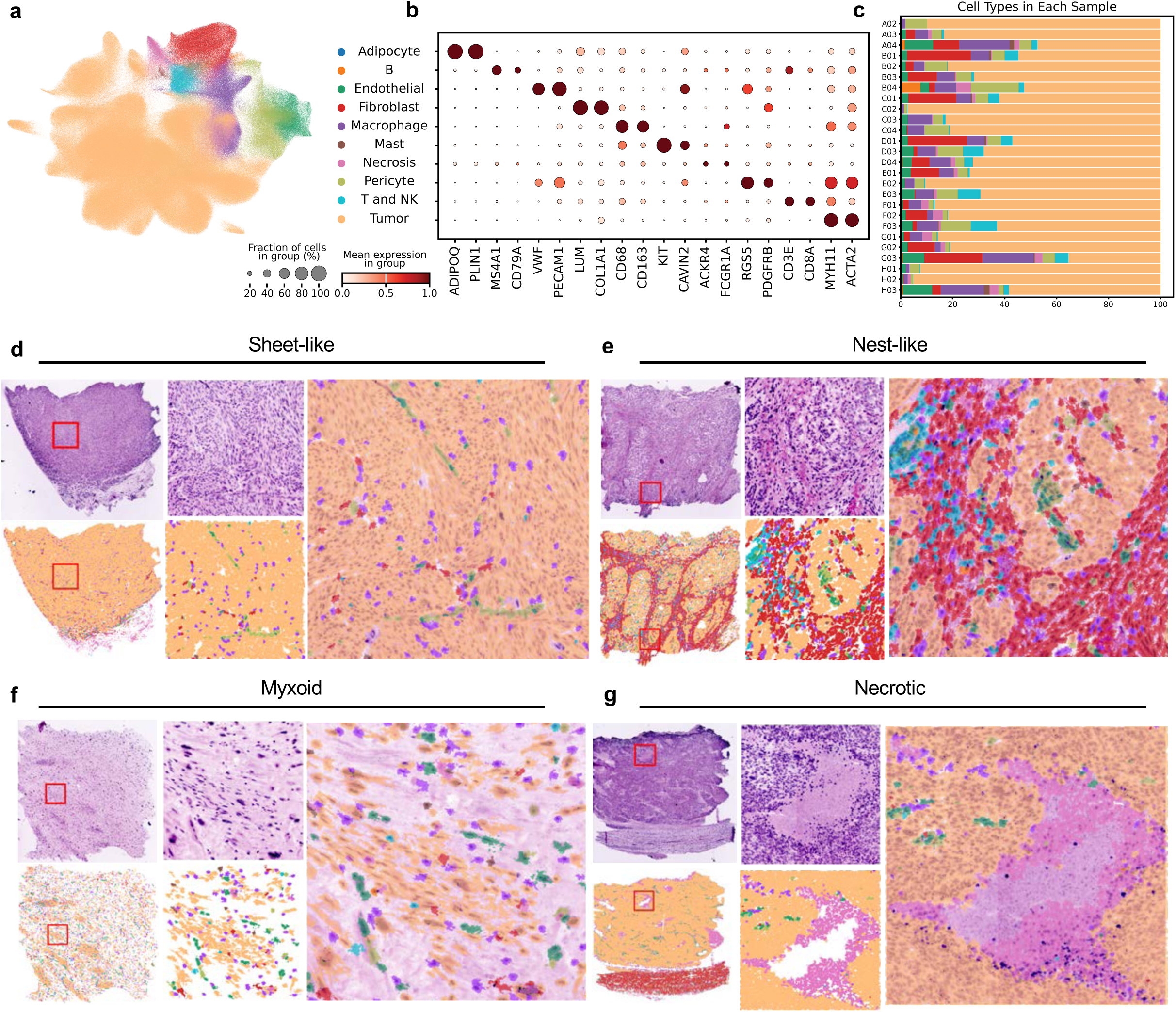
Broad spatial morphologies of ULMS tumors. **a,** Batch-corrected UMAP of all segmented cells from G4X spatial transcriptomic tissue sections, colored by annotated cell type. **b,** Dot plot showing mean expression of selected markers across annotated cell types. **c,** Proportion of each cell type across patient tissue sections. **d-g,** Representative fluorescent H&E images and corresponding spatial transcriptomic annotation and segmentation. Fluorescent H&E (top) and cell type annotations (bottom), both at low power (left) and higher magnification (middle), and merged overlay showing spatial localization of cell populations (right). Representative images include patterns showing sheet-like morphology (**d**), nest-like morphology (**e**), myxoid pattern (**f**), and necrotic areas (**g**).

Based on the morphology of these tumors on histology alone, there were variations across samples that were consistent with previously described histological variations and subtypes in ULMS (**Extended Data Fig. 17**). A sheet-like formation comprised predominantly of tumor cells with interspersed blood vessels was the most common histological pattern across patient samples (**Fig. 4d and Extended Data Fig. 17**). Additionally, some samples had tumor cells growing in nest-like groups within a fibrotic extracellular matrix (**Fig. 4e and Extended Data Fig. 17**). An additional morphology was a myxoid pattern, which correlated with the samples clinically diagnosed as myxoid leiomyosarcoma, with abundant stroma and scattered tumor cells and other cell types (**Fig. 4f and Extended Data Fig. 17**). Finally, several tumor samples were pockmarked with acellular areas of necrosis surrounded by tumor cells (**Fig. 4g and Extended Data Fig. 17**).

### Spatial morphologies of tumor states complement the scRNA-seq derived transcriptomic phenotypes

We then turned to further dissecting the spatial tumor states, by subsetting, re-integrating, and clustering the tumor cells. Analogous to the tumor states identified using scRNA-seq, the spatial tumor subclusters broadly separated on the basis of SMC markers, with some tumor cell states expressing higher levels of SMC markers (ESR1 PGR AR Tumor, CHI3L1-high SMC-like Tumor, CHI3L1-low SMC-like Tumor, IFN Signaling Tumor) and others with lower expression of SMC markers (Ischemic Tumor, COL1A1 POSTN Tumor, and SDC1 PDGFRB Tumor) (**Fig. 5a, b, and Extended Data Fig. 18, 19a, b**). There were also several clusters with relatively higher expression of proliferation markers and varying levels of canonical SMC markers (Cycling IFN Signaling Tumor, Cycling SMC-low Tumor, and Cycling COL1A1 POSTN Tumor cells).

**Figure 5.**
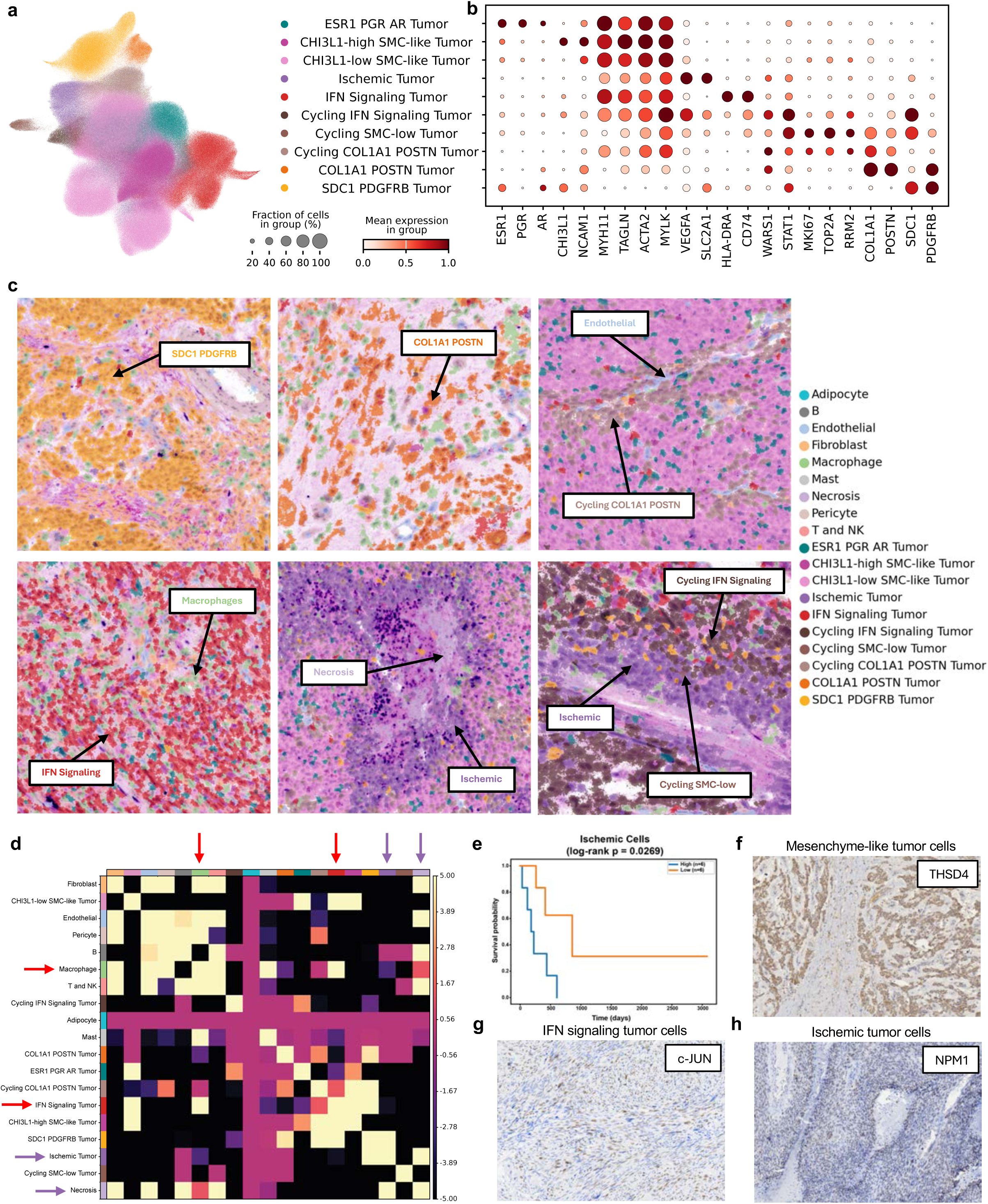
Spatial organization patterns of the dominant ULMS tumor cell states. **a,** Batch-corrected UMAP of spatially defined tumor cell states from G4X sections, colored by annotated ULMS tumor cell states. Colors broadly parallel their inferred corresponding cell identities on scRNA-seq (Fig. 2). **b,** Dot plot showing mean expression of selected markers for each spatial tumor state. **c,** Representative spatial regions highlighting several spatial tumor cell states identified. **d,** Heat map summarizing neighborhood enrichment analysis across all samples for all tumor cell states. Red arrows highlight the association between macrophages and IFN Signaling Tumor cells. Purple arrows highlight the association between necrosis and Ischemic Tumor cells. **e,** Kaplan-Meier curve plotting the outcomes of patients with high and low enrichment of the ischemic cell signature using bulk RNA sequencing data of ULMS samples from TCGA. **f-h,** Representative IHC sections showing positive staining for each predominant tumor state identified, namely THSD4 for Mesenchyme-like tumor cells **(e)**, c-JUN for IFN signaling tumor cells **(f)**, and NPM1 for Ischemic tumor cells **(g)**.

Although the G4X assay was comprised of a restricted panel of genes, we were able to broadly map the spatial tumor cell states to the scRNA-seq-defined tumor cell states using a combination of canonical markers and pathway enrichment analysis that overlapped across both assays, particularly for the three predominant scRNA-seq-defined tumor cell states. First, the COL1A1 POSTN and SDC1 PDGFRB Tumor cell states identified spatially both had enrichment of markers and pathways associated with collagen degradation and ECM remodeling, which aligned with the Mesenchyme-like tumor cell state defined on scRNA-seq (**Fig. 5a, b, and Extended Data Fig. 19a**). Interestingly, the spatial morphologies of these cells complemented their transcriptomic phenotypes: the SDC1 PDGFRB cells formed nests of cells surrounded by fibrotic areas, the COL1A1 POSTN cells were found scattered throughout tumor sections, particularly in myxoid tumor samples, and the Cycling COL1A1 POSTN tumor cells frequently localized around blood vessels (**Fig. 5c, and Extended Data Fig. 18, 19b**). These spatial phenotypes highlight the mesenchyme-like features of these cells and likely reflect the high plasticity and potential for increased mobility and invasiveness of these cells.

Second, the spatial IFN Signaling Tumor cells had high expression of *HLA-DRA*, *CD74*, *WARS1* and *STAT1*, suggestive of active interferon signaling, which aligned with the IFN signaling tumor cell state identified on scRNA-seq (**Fig. 5a, b, and Extended Data Fig. 19a**). These IFN Signaling Tumor cells were found scattered throughout tumor sections, at times forming sheet-like patterns (**Fig. 5c, and Extended Data Fig. 18, 19b**). On scRNA-seq, the IFN signaling tumor cells were found to have high enrichment of both NFKB and AP-1 signaling, along with high expression of *CD47* that complemented the elevated *SIRPA* among macrophages and OGCs, suggestive of these cells employing anti-phagocytic mechanisms to promote an immunosuppressive microenvironment. To further validate this mechanism, we computed spatial neighborhoods across all spatial samples. This analysis revealed proximity of macrophages to IFN Signaling Tumor cells, further supporting their previously inferred immunosuppressive interactions (**Fig. 5c, d, and Extended Data Fig. 20**).

Third, the spatial Ischemic Tumor cells had high expression of *VEGFA* and *SLC2A1* and were consistently located around areas of necrosis, presumably in the ischemic penumbra, which aligned with the Ischemic tumor cell state defined on scRNA-seq. These cells were enriched for apoptosis and programmed cell death pathways, which were also enriched in the cycling tumor cell states (**Fig. 5a-c, and Extended Data Fig. 19a**). Furthermore, the Ischemic tumor cells were highly associated with necrotic regions on neighborhood enrichment analysis (**Fig. 5d, and Extended Data Fig. 20**). These findings were suggestive of highly proliferative areas encompassing necrotic regions due to high cell turnover that were surrounded by Ischemic Tumor cells in the penumbra. Necrosis is one of the three main diagnostic criteria for ULMS,^80,81^ and both high degrees of necrosis and the immunohistochemical presence of MYC have been shown to confer worse prognoses.^57,82^ To investigate whether the transcriptomically defined Ischemic Tumor cells identified in this study correlate with the previously described clinical-pathological findings, we analyzed how patient outcomes differed based on the enrichment of this tumor cell state in bulk RNA sequencing data from TCGA. Patients with high enrichment of the Ischemic Tumor cells had significantly worse outcomes (**Fig. 5e; Extended Data Fig. 21 a-d**). This concordance between our transcriptomically defined cell state and previously established histopathological features supports this tumor state classification as biologically meaningful.

To investigate whether these bioinformatically defined tumor states were representative of biologically relevant phenotypes, IHC was performed for markers demonstrative of each major tumor state, using matched ULMS tumor samples. THSD4 (Thrombospondin Type 1 Domain Containing 4), an ECM regulatory protein highly expressed in the Mesenchyme-like tumor cells, was highly expressed in tumor cells forming nest-like morphologies and scattered throughout tumor cells across several samples (**Fig. 5f, Extended Data Fig. 22a**). JUN localized to tumor cells scattered throughout sections and in sheet-like regions across several samples (**Fig. 5g, Extended Data Fig. 22b**). NPM1 (Nucleophosmin 1), a transcriptional target of MYC, was expressed in significantly high levels around areas of necrosis, consistent with the spatial localization of the Ischemic tumor cells (**Fig. 5h, Extended Data Fig. 22c-d**).

### Spatial localization patterns of hormonal tumor cells support their stem cell-like transcriptomic profile

Given their stem-like phenotype on scRNA-seq, we turned to further profiling the spatial characteristics of the ESR1+ cells and the AR+/PR+ cells. Spatially, the ESR1 PGR AR Tumor cells had high expression of hormonal receptors that were representative of two hormonal clusters identified on scRNA-seq (**Fig. 5a, b, and 6a, b**). While standard hormone receptor immunohistochemistry provides spatial localization, it should be noted that the ESR PGR AR Tumor cluster is defined not only by hormone receptor expression, but also by the expression of other transcripts in the panel, such as CCND1 and CD44, as well as by its spatial neighbors. This is further supported by Moran’s I analysis demonstrating high dispersion of *ESR1*, *PGR*, and *AR* (**Extended Data Fig. 23a, b**). This spatial cluster had high expression of SMC markers, corresponding more to the gene expression profile of AR+/PGR+ cells (**Fig. 6b**). It is likely that this ESR1 PGR AR Tumor cluster represents two cell states that were identified as one cluster spatially due to the lack of additional distinctive markers to differentiate the two populations with more granularity.

**Figure 6.**
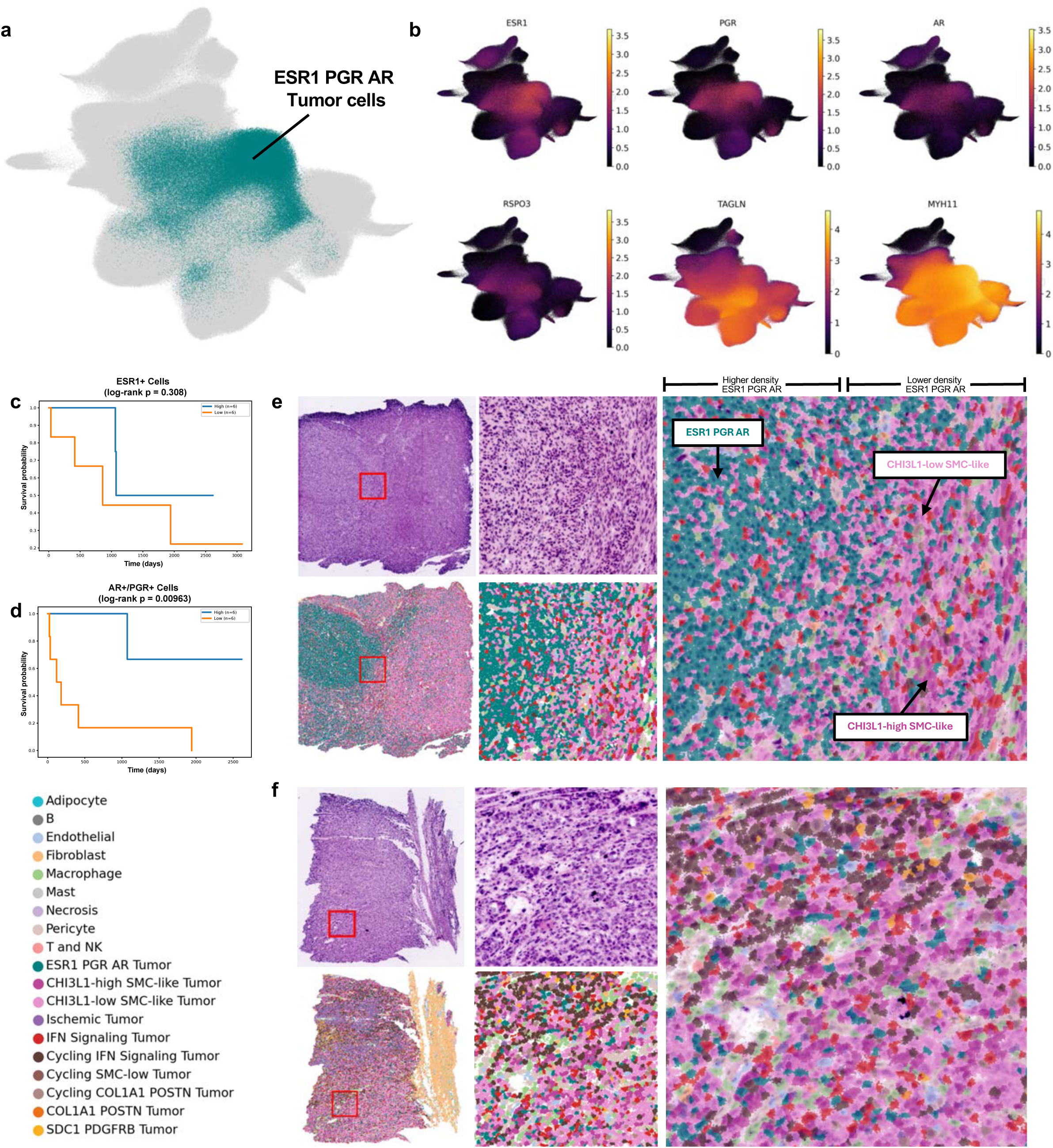
Spatial distribution of ESR1 PGR AR tumor cells. **a,** Batch-corrected UMAP of spatially defined tumor cells from all G4X sections, with the ESR1 PGR AR cluster colored. **b,** Markers representative of the ESR1 PGR AR cluster plotted on the batch-corrected UMAP. **c-d,** Kaplan-Meier curves plotting the outcomes of patients with ULMS with bulk RNA sequencing data from TCGA, with high and low enrichment of ESR1+ tumor signature and AR+/PGR+ tumor signature. **e-f**, Representative spatial regions highlighting the ESR1 PGR AR cell type. Fluorescent H&E (top) and cell type annotations (bottom), both at low power (left) and higher magnification (middle), and merged overlay showing spatial localization of cell populations (right).

To additionally validate that the hormonal cells were two distinct phenotypic clusters with differing biological significance, and to further substantiate these clusters as biologically meaningful states that are consistent with previously established literature, we analyzed patient outcomes based on the enrichment of the signatures from these two clusters in bulk RNA sequencing data from The Cancer Genome Atlas (TCGA). Prior histological studies have demonstrated improved prognosis in patients with ULMS with high expression of androgen and progesterone receptors.^83–86^ Consistent with these previous studies, in our analysis, patient survival was not significantly correlated with the ESR1+ signature, whereas high enrichment of AR+/PGR+ cells was associated with a significant survival benefit (**Fig. 6c, d, Extended Data Fig. 22, and Supplementary Table 7**). The concordance of the tumor states described in this study with previously described histopathological and clinical findings further corroborates the tumor states described here as biologically and clinically relevant.

We next sought to understand the spatial arrangement of hormone receptor-positive cells by exploring their cellular organization (**Fig. 6e, f, Extended Data Fig. 18**). There were some sections where the ESR1 PGR AR Tumor cells co-localized in high density (**Fig. 6e, Extended Data Fig. 18**); even within these sections, there were gradients of these hormonal cells, with areas of high density with adjacent areas of lower density of these ESR1 PGR AR Tumor cells. Within these gradients, the areas with lower ESR1 PGR AR tumor cells showed a diversity of other tumor cell states, particularly CHI3L1-high and CHI3L1-low SMC-like Tumor cells. The ESR1 PGR AR tumor cells also appeared near various cycling tumor cell states (**Fig. 6f, Extended Data Fig. 18**). Given these findings, it is possible that the areas where the ESR1 PGR AR Tumor cells congregate in high density may be leading edge areas of active tumor growth and proliferation, where stem-like cells differentiate into other tumor cell states. In fact, on neighborhood analyses, the ESR1 PGR AR Tumor cells most strongly colocalized with CHI3L1-low SMC-like Tumor cells, followed by CHI3L1-high SMC-like Tumor cells and the Cycling IFN Signaling Tumor cells (**Fig. 5d, Extended Data Fig. 20**). In sum, these findings support the hypothesis that hormonal receptor-expressing cells play a role in ULMS as niche and stem-like cells that promote the survival and growth of these tumors.

### Drug sensitivity modeling predicts alternative targeted candidates for tumor cell states predicted to be resistant to first-line therapies

Most patients with ULMS do not respond well to first-line therapies, a phenotype that is well represented among the recurrent and metastatic cohort analyzed in this study. It is unclear whether the transcriptomic heterogenous ULMS tumor cell states may be predictive of varying treatment sensitivities, and if so, whether phenotypes less sensitive to first-line chemotherapies would demonstrate superior vulnerabilities to other targeted therapy options. To understand the drug sensitivities of the different tumor cell states, we applied the single-cell integration and drug response prediction (scIDUC) algorithm to all tumor cells from our scRNA-seq analysis. By correlating bulk RNA sequencing data from cancer cell lines with high-throughput cancer cell line drug screening data, the scIDUC algorithm predicts the likelihood of response (depicted as nAUCs) for each compound available in the CTRPv2 database per cell.^87^ In contrast to most other drug prediction algorithms, scIDUC is thus based on experimental drug screening data. Certain compounds were omitted from further analysis due to the absence of a statistically significant gene expression signature associated with their efficacy. A lower nAUC for a given drug indicates a lower dose necessary for cell death, implying higher vulnerability to that drug.

Overall, the drug response patterns across the different tumor cell states were very heterogenous (**Fig. 7a and b; Supplementary Table 8**). The ESR1+ cells and the AR+/PGR+ cells were predicted to be sensitive to most drugs in the database, including first-line ULMS chemotherapies gemcitabine, docetaxel, and doxorubicin. Given that almost all samples in this dataset were from patients who had undergone a variety of different chemotherapy treatments prior to sample collection, it is possible that these hormone receptor-expressing cells were relatively lower in abundance due to their increased sensitivity. ATRX+/DAXX-tumor cells were also predicted to be highly sensitive to many drugs. All other tumor cell states were predicted to be less sensitive to most drugs, consistent with the aggressive clinical phenotype of ULMS tumors.

**Figure 7.**
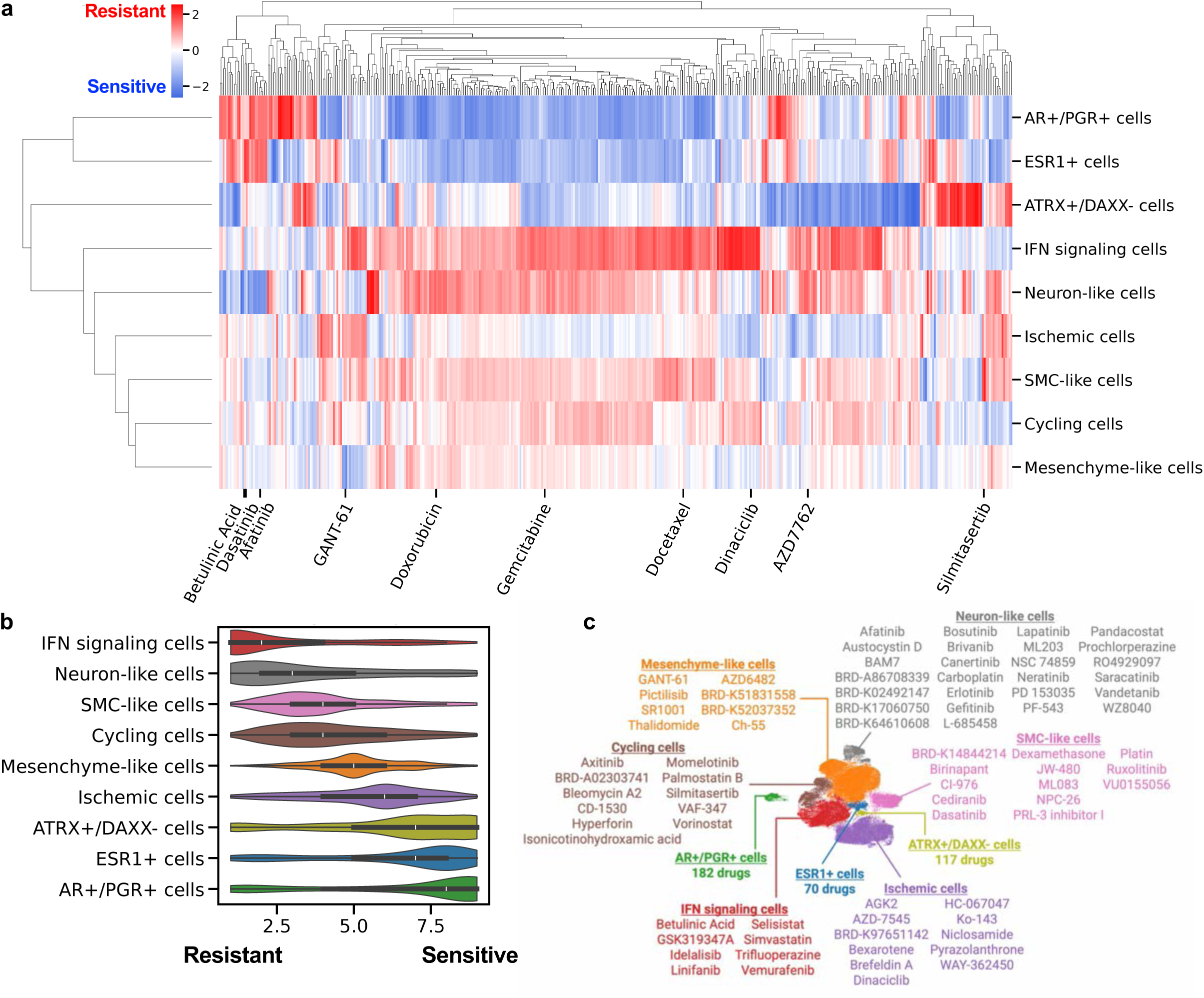
Drug sensitivity predictions using the scIDUC algorithm. **a**, Heat map depicting Z-scaled predicted nAUCs of each tumor cell state across all drugs, spanning a spectrum of highest (blue) to lowest (red) sensitivity. **b**, Box plot depicting the mean rank of drugs within each tumor cell state, with Rank 9 having the lowest nAUC (most sensitive) and Rank 1 having the highest nAUC (most resistant) for each drug. **c**, List of drugs ranked 9 (most sensitive) within each tumor cell state. All drugs and their ranks per cluster are listed in detail in **Supplementary Table 8**.

Even though the more chemoresistant tumor cell states appeared resilient to most drugs tested, each of these tumor cell states was predicted to be uniquely sensitive to a small subset of drugs. To identify the drugs with the highest predicted efficacy for each tumor cell state, the average nAUC of all cells belonging to each cluster was computed, and each drug was matched to the cluster with the lowest nAUC (i.e. most sensitivity) for that drug (**Fig. 7c; Supplementary Table 8**). As expected, the ESR1+ and AR+/PGR+ tumor cells had the greatest number of drugs predicted to be effective against them.

All other tumor cell states had lower numbers of uniquely efficacious drugs predicted. Some of these drugs had known mechanisms that targeted susceptibilities unique to each cluster. For example, Dinaciclib, a CDK9 inhibitor, was predicted to be highly efficacious against the Ischemic tumor cells. Prior work has shown that Dinaciclib is particularly efficacious against other MYC driven cancers, along with pre-clinical work suggestive of efficacy in osteosarcoma and soft tissue sarcoma models.^88–90^ Betulinic acid, predicted to have the highest efficacy against the IFN signaling cells, is a naturally occurring compound that has been shown to reduce interferon-stimulated genes and TNF-mediated NFKB stimulation,^91–93^ prominent signatures in these IFN signaling cells. Among the Mesenchyme-like cells, the drug predicted to be most highly efficacious was GANT-61, a GLI inhibitor that has been shown to inhibit epithelial-to-mesenchymal and myofibroblastic transition, with pre-clinical efficacy demonstrated in other sarcomas.^94–96^ These results offer predictions for alternative therapy options for tumor states with inferred resistance to first-line therapies.

## Discussion

We present the first comprehensive transcriptomic single-cell and spatial atlas of ULMS, profiling 23 recurrent, metastatic ULMS tumors from 18 patients. We define multiple tumor cell states that are present across multiple scRNA-seq samples and identify a spatial correlate for each tumor cell state, specifically highlighting 1) hormone receptor-positive stem-like cells scattered throughout the tumor parenchyma, 2) inflammatory tumor cells characterized by interferon signaling, 3) ischemic tumor cells surrounding areas of necrosis, and 4) invasive mesenchyme-like cells growing in nests or surrounding blood vessels. The unique transcriptomic signatures of these clusters speak to the complex biology driving ULMS growth and progression.

The tumor cell states identified in this atlas overlap with prior classifications of leiomyosarcomas based on bulk transcriptomics and genomics, which we summarize briefly here. Beck et al. describe three clusters: group I with enrichment of muscle contraction and actin cytoskeleton related genes; group II enriched for genes associated with protein metabolism, cell proliferation, and organ development; and group III related to organ and system development, extracellular proteins, and protein synthesis.^23^ Guo et al. also describe three subtypes: subtype I LMS, which shows upregulation of the transcription factor *MYOCD*, which is known to be amplified in many ULMS patient samples;^19,25^ subtype II LMS, which shows upregulation of translation; and subtype III LMS, which shows expression of metabolic and transcriptional pathways.^24^ Chudasama et al. also define three subtypes: subgroup 1 that has high enrichment of platelet degranulation, complement activation, and metabolism; subgroup 2 that had high enrichment of muscle-related genes, including *ARL4C*; and subgroup 3, also with muscle-related gene expression and cell-cell signaling, with high *CASQ2* and *LMOD1*.^21^ Hemming et al. identify three LMS subtypes: conventional (smooth muscle-like), inflammatory (upregulation of immune-related genes), and uterogenic (composed mostly of uterine-derived tumors).^25^ Most recently, Anderson et al. divide LMS into three developmental subtypes: subtype 1 (mixed dedifferentiated tumors), subtype 2 (LMS of vascular origin), and subtype 3 (LMS mostly of gynecological origin).^26^

As described in the 2022 consensus summary by Burns, Jones, and Huang, these prior studies of leiomyosarcomas all identify three major subtypes using bulk transcriptomic and genomic approaches, with shared features across their classification systems that also overlap with the tumor states in our study (**Extended Data Fig. 15a-d**).^97^ For example, one consistently described feature across studies is a subtype with enrichment of myogenic markers (such as Beck group I, Guo subtype I, Chudasama subgroups 2 and 3, Hemming cLMS, and Anderson subtype 2), which overlaps with the tumor cell states expressing high SMC markers in our study, including AR+/PGR+ cells, ESR1+ cells, SMC-like cells, IFN signaling cells, and cycling cells. Another feature identified in subtypes across studies is high translational and ribosomal activity (such as Beck group III and Guo subtype II), which correlate to the ischemic, cycling, and IFN signaling cell states in this study. Tumor cells with high metabolism is another shared feature across studies (such as Beck group II, Guo subtype III, and Chudasama subgroup 1), which overlap with the IFN signaling cells in our study. However, the previously described bulk transcriptomic signatures likely derive from varying proportions of more granular single-cell states within the samples, and therefore the tumor states identified in our study are likely overlapping across multiple prior classifications.

In this single-cell study, differentiation status relates to developmental potency, not histological phenotype as commonly referred to in pathological specimens. As an example, AR+/PGR+ stem-like cells with relatively higher predicted potency have smooth muscle-like features. By contrast, Mesenchyme-like and Neuron-like cells with lower predicted potency do not resemble smooth muscle cells and would be histologically characterized as “dedifferentiated.” The contradiction in terms between potency (developmental potential) and differentiation (tissue resemblance) further highlights an opportunity for single-cell investigation of sarcomas. Our data suggest that while ESR1+ stem-like cells may have the capacity for tumor renewal, histologically dedifferentiated ULMS cells may not have the developmental potency to return to a smooth-muscle-like state; this remains to be experimentally validated.

Based on canonical uterine stem cell markers such as CD44^39^ and CytoTRACE 2 results, we theorize that ESR1+ cells are stem-like sarcoma cells that help to regenerate and renew the tumor. The developmental lineage of these cells remains unclear. ESR1+ cells show the highest expression of the Wnt/beta-catenin pathway, which is known to support endometrial-mesenchymal stem-like cells and fibroid stem cells.^37,38,40^ However, fibroid stem cells typically express less estrogen receptor than the surrounding cells and rely on paracrine signaling from the stem cell niche.^37–39^ Expression of ESR1 suggests that ULMS stem-like cells have switched to an autocrine mechanism to support tumor growth. Given the genetic similarity among ESR1+ ULMS stem-like cells, endometrial-mesenchymal stem-like cells, and fibroid stem cells, future studies should investigate whether these cells share a common lineage.

Drug prediction based on correlation with *in vitro* drug screening data shows that each tumor cell state may have unique therapeutic vulnerabilities, with inferred drug sensitivities varying widely among the different states. For example, the ESR1+ cells and AR+/PGR+ cells were predicted to be sensitive to most drugs tested, including current first-line chemotherapy agents; the broadly increased susceptibility of these cells to chemotherapy may play a role in the improved survival observed in patients whose tumors are enriched with the AR+/PGR+ cell phenotype. The maximal sensitivity of ESR1+ cells and AR+/PGR+ cells to current first-line chemotherapy regimens suggests that these options should be maximally exhausted in patients with a high degree of positive hormone receptor staining. However, our inferences suggest that alternative agents may be necessary for the other tumor cell states present in the tumor.

This study has several limitations. Given that most patients had undergone several rounds of chemotherapy, radiation, and surgery prior to sample collection, it is likely that the biological mechanisms identified in this study are influenced by prior treatments. This study was performed on a cohort of treatment-resistant and recurrent ULMS tumors, and it is unclear whether the findings from this study are also reflective of primary ULMS tumor biology. Future studies of primary ULMS tumors are needed to bridge this gap. Another limitation of this study is in the lack of further understanding of the exact nature of the stem-like hormone receptor expressing cells. Specifically, although CytoTRACE 2 computes scores reflective of absolute differentiation potential per cell, it is unknown whether the less differentiated cells were previously mature cells that later dedifferentiated or if they are stem-like cells that never differentiated. Further exploration of these cells and their exact origins and differentiation characteristics are needed. Furthermore, correlation between scRNA-seq and spatial transcriptomic data was challenging largely due to the limited number of genes available in the G4X spatial transcriptomic panel. The broader cell types and tumor states identified with scRNA-seq could be correlated to tumor clusters with high and low expression of canonical SMC markers in the G4X spatial data; however, finer resolution correlation between the two platforms was more challenging due to technical differences between the assays and sampling bias. Though we were able to match most tumor cell states using a combination of canonical markers and pathway analysis that correlated across platforms, this led to some scRNA-seq defined tumor states not having clean delineations in the G4X spatial data, such as the two hormone receptor expressing clusters identified with scRNA-seq identified as one spatial cluster. While IHC supports the definition of most tumor states identified with scRNA-seq, additional spatial studies are needed using targeted or more comprehensive gene panels for further validation. Finally, though the scIDUC drug predictions were based on integration of bulk RNA sequencing, scRNA-seq, and cancer cell line testing data that is conceptually robust, these are only hypothesis-generating drug predictions and need rigorous experimental validation. Currently available models for ULMS are limited and have been shown to be poorly representative of primary tumor states. Therefore, there is an urgent need to develop additional experimental models of ULMS that fully recapitulate the complex tumor states and heterogenous microenvironment of these tumors. Future studies are needed to develop such experimental ULMS models, including cell lines representative of the different ULMS tumor states and organoids that recapitulate the complex ULMS TME, to further validate the drug sensitivity predictions in this study. Without such validation, the validity and applicability of these inferences remain limited.

In summary, we present a comprehensive transcriptomic atlas of ULMS with spatial granularity, defining tumor states of various cellular potencies that interact to produce a recurrent metastatic tumor. ESR1+ stem-like cells are likely sensitive to current regimens, while other less potent but more histologically dedifferentiated cell types are less sensitive to standard chemotherapy regimens. These findings have implications for rational design of systemic and regional therapies for ULMS.

## Methods

### Patient Tissue Collection and Processing

This study was performed under IRB protocol IRB-65893 at Stanford University. Patients undergoing surgery at Stanford Hospital were approached preoperatively, and voluntary, written, informed consent was obtained from every patient. Where applicable, some patients were separately consented by an independent study team for cytoreduction and HIPEC under a phase 2 trial,^98^ which was distinct and independently approved by a different review board from our study. This study is not a translational aim from that trial. At the time of tumor resection, a sample of the resected tumor was collected. Specimens were transported on ice and immediately dissociated into single-cell suspensions using the standard protocol for the gentleMACS Octo Dissociator (Miltenyi, #130-134-029 and #130-095-929). If necessary, red blood cell lysis was performed for 10 minutes at room temperature using RBC lysis buffer (Roche, #11814389001). The samples were then either cryopreserved as live cell suspensions or directly carried through to library preparation and sequencing.

### ScRNA-seq Library Preparation and Sequencing

Given the rarity of ULMS, the tumor samples used in this study were collected over the course of several years, over which time the exact cell preparation, library preparation methodologies, and sequencing platforms evolved. All library preparations were performed using 10x Genomics protocols, and all samples underwent single-cell RNA sequencing. If cryopreserved as live cell suspensions, cells were thawed, dead cell removal was performed (Stem Cell Technologies, #17899), and cells were counted. Once optimal concentration was achieved, we proceeded with library preparation using standard 10x Genomics protocols per chemistry. The chemistry used for each sample is listed in **Supplementary Table 9**. Samples were sequenced across three platforms: 1) at the Stanford Functional Genomics Facility on the Illumina NovaSeq 6000, 2) an Illumina Novaseq sequencer through Retro Biosciences, or 3) the Singular Genomics G4 sequencer after undergoing library conversion for sequencer compatibility. FASTQ files from all batches were aligned independently using 10x Genomics Cell Ranger v8.0.1 to the reference genome GRCh38-2024-A.^99^

### Preprocessing and Quality Control

Ambient RNA removal was performed using cellbender 0.3.0.^100^ Learning rate and maximum epochs were modified to achieve appropriate convergence. Filtered h5 files were then imported into python as anndata objects.^101,102^ Preprocessing was performed using the scanpy framework on each batch^103^ with a minimum of 100 genes per cell and the maximum percentage of mitochondrial transcripts per cell set to 50%. These metrics were chosen based on visual inspection of batch-level quality control plots, with the goal of initial lenient cell filtering. Due to overrepresentation of TCR and BCR genes in one sample from concomitant TCR sequencing, these genes were removed from all batches. Doublet removal was performed on each batch using DoubletDetection^104^ with n_iters=20, p_thresh=1e-16, and voter_thresh=0.5.

### Integration, Clustering, and Differential Gene Expression

The top 2000 highly variable genes were identified using the seurat_v3 method in scanpy. Integration was performed using scVI with the sequencing batch and patient sample as categorical covariate keys.^32,105,106^ RBCs and a low-quality cell cluster were removed following initial integration. The high-quality cells were then reintegrated with scVI for the default number of epochs, and Leiden clustering was performed on the scVI embedding.^107^ Differential expression analysis was performed using scVI and scanpy.

### Benchmarking Integration

We used scIB-metrics to benchmark the integration of all cells after quality control.^33,34^ This package uses multiple metrics to compare bioconservation and batch correction of different integration methods. After quality control, data were normalized and log transformed. The top 2000 highly variable genes were identified using the seurat_v3 method in scanpy. PCA was computed using 50 principal components; this was the “Unintegrated” representation. The python implementation of Harmony^108^ was then run on these principle components using batch and sample as the integration keys. This was compared to the scVI representation using default parameters of the benchmark() function.

### Subset Analyses

When further analyzing subsets of cells, such as subsets of lymphoid cells, myeloid cells, and tumor cells, each of these cell types were independently subsetted, filtering out all other cell types in the downstream analysis. For example, when further analyzing tumor cell sub-states, only the tumor cells were included, filtering out all other cell types. Before integrating each subset, the top 2000 highly variable genes were independently chosen for each subset using the seurat_v3 method in scanpy, filtering low-abundance genes and increasing the span as needed. The subsetted cells were then integrated with scVI separately and manually annotated by Leiden cluster, using the top differentially expressed genes per cluster and comparison to publicly available data. For the tumor cell subset, Louvain clustering was also performed to further assess the robustness of the Leiden clusters.^107^

### Downstream Analysis

Copy number inference was performed using the python implementation of CNV. This method has been previously described.^35,36^ In brief, a gene order file was constructed from the GRCh38-2024-A reference genome gtf file (from 10X Genomics) using the gtf_to_position_file.py script provided by the InferCNV authors (https://github.com/broadinstitute/inferCNV/tree/master/scripts). The gene order file was mapped to the available variables using Ensembl IDs. Gene expression was then averaged over a running window of 500 genes, compared to that of a reference cell type, and denoised, as previously described (reference). A window of 500 genes was chosen to better highlight large-scale CNV aberrations in ULMS. Immune cells (B cells, myeloid cells, T and NK cells, mast cells, plasma cells, and plasmacytoid dendritic cells) were used as the reference cell type. Inferred CNV was computed on all batches together.^35,36^

We used the default method in Decoupler to perform transcription factor inference; this method has been previously described in detail.^55^ The tumor subset anndata object was normalized, log transformed, and scaled to mean of zero and variance of one, with a maximum expression cutoff of 10. Briefly, the CollecTRI database of transcription factor regulons was downloaded from Omnipath.^109,54^ Each regulon contains a transcription factor and its putative dependent genes, with weights showing the direction and strength of the interaction. For each gene, a univariate linear model was constructed, with gene expression as the dependent variable and the weight of that gene in the regulon as the independent variable. An enrichment score for the transcription factor of interest was then calculated as the t-value of the slope coefficient of the linear model. The enrichment scores for each tumor cluster compared to all other clusters were ranked using Decoupler’s implementation of the Wilcoxon test in rankby_group() and plotted using Scanpy’s matrixplot(). The same univariate linear model was used for analysis of upregulated molecular pathways in the tumor clusters using Reactome gene sets, again downloaded from Omnipath.^70,109^ For Reactome pathway analysis, manually selected pathways with significant enrichment per cluster were plotted using Scanpy’s matrixplot().

Cellular potencies for each tumor cell were inferred using the python package for CytoTRACE 2.^43^ Given the rarity of these tumors and to avoid excluding any rare tumor cell phenotypes, raw counts from all tumor cells were input together, without filtering for gene counts. As such, the pre-KNN CytoTRACE 2 scores were used for downstream analysis and interpretation. Cell cycle scores were computed using the score_genes_cell_cycle function in scanpy, using a publicly available list of cell cycle genes.^110^

After standard preprocessing and calculating the centroid of each of the tumor clusters, the SCimilarity reference database was searched for the top 10,000 most similar cells.^48^ The tissue of origin and cell type of these cells were plotted.

Cell-to-cell interactions were inferred using the CellphoneDB tool, using the statistical analysis method (method 2) to predict significant interactions. Results were visualized using the functions in ktplotspy.

### Cluster Annotations

For annotating broad cell types, such as for all cells and for the myeloid and lymphoid subpopulations, expression patterns of canonical markers were visualized on UMAP projections. The optimal clustering resolution was chosen based on assessment of feature plots, along with evaluating top differentially expressed genes for each cluster over a spectrum of resolutions. For annotating tumor cell states, after similarly evaluating a spectrum of resolutions, the final clustering resolution was selected based on clusters that had differentially expressed genes that were suggestive of biologically meaningful and coherent programs. This was chosen based on literature review of top differentially expressed genes, along with review of the results from the downstream analyses described above, including CNV inference, transcription factor predictions, differentiation potentials, and SCimilarity.

### 3SEQ Correlation Analysis

3SEQ data was downloaded from GSE45510. Each tumor state’s signature was identified by filtering only for statistically significant genes (p <0.05) that were also present in the 3SEQ dataset, and subsetting the top 50 differentially expressed genes by either logFC or scanpy score. The Gseapy package was used to run ssGSEA,^111^ and normalized enrichment scores were plotted per tumor state signature for each sample.

### Singular G4X Tissue Preparation and Sequencing

Samples were prepared from formalin-fixed, paraffin-embedded (FFPE) patient blocks according to the manufacturer’s protocol.^78^ Briefly, FFPE blocks were acquired from the Department of Pathology archive and freshly sectioned at a thickness of 5 µm onto custom gel pads. Regions of Interest (ROIs) were selected based on hematoxylin and eosin (H&E) stains previously performed by the Department of Pathology, with areas of high tumor enrichment annotated by a pathologist. ROIs were manually punched out of the gel pad and transferred onto functionalized glass slides. After deparaffinization, rehydration, and antigen retrieval, hybridization with the Singular G4X breast cancer panel (309 genes) was performed. Ligation of the probes and rolling circle amplification were then performed *in situ*. The slide was conditioned, framed, and loaded onto the G4X instrument. Fluorescent H&E and spatial sequencing were performed on the G4X instrument.

### Singular G4X Segmentation and Preprocessing

We used the onboard Cellpose segmentation as the initial nuclei-based segmentation state for Proseg segmentation after nonspecific probes were removed.^79,112^ Data were explored in QuPath 0.5.1 and imported into python using Anndata and Spatialdata containers.^102,113^ Quality control was performed on each tissue section using a minimum of 20 counts and 5 genes. Cells at the 0.5% and 99.5% extremes for Proseg segmentation area were excluded. Spatial outliers were removed individually for each section by computing a spatial nearest neighbors graph for *n* neighbors and removing cells with a minimum Euclidean distance of *x* standard deviations, with *n* and *x* customized for each tissue section.

### Singular G4X Integration, Clustering, and Differential Gene Expression

After normalization and log transformation, all cells passing quality control were integrated using an unsupervised resolVI model with each tissue section as the batch key.^114^ The model was trained for 100 epochs. Data were scaled to a mean of zero and a variance of one, and maximum gene expression was clipped at 10. Leiden clustering was performed, and differentially expressed genes were computed using the Scanpy implementation of the Wilcoxon test. Coarse cell types were annotated at resolution 0.4 based on expression of canonical markers. The resolVI embedding and coarse cell type annotations were then used as the input to the scVIVA model.^115^ This model was constructed using the k=20 nearest neighbors of each cell, with the patient selected as the predominant source of batch effect. The model was trained for a maximum of 600 epochs with early stopping, a batch size of 512, and a learning rate of 0.0005. Leiden clustering was then performed on the scVIVA embedding, and differentially expressed genes were again computed using the Scanpy implementation of the Wilcoxon test. After initial annotation of Leiden clusters at resolution 0.5, subclustering of tumor, vascular, fibroblast, and immune populations was performed, and each of these subclusters was manually annotated based on expression of canonical markers as well as differentially expressed genes. Normalized and corrected counts generated by the resolVI model were used extensively during this iterative manual annotation process in an attempt to correct for transcript misassignment.

### Correlation of G4X and scRNA-seq Tumor Clusters

G4X and scRNA-seq tumor clusters were correlated based on the presence or absence of smooth muscle markers, top differentially expressed genes, and pathway enrichment scores. Pathway enrichment scores for Reactome pathways were calculated using Decoupler’s univariate linear model in the same way as described above for scRNA-seq tumor states, and the top pathways per cluster were ranked using the Wilcoxon test in Decoupler’s rankby_group().^55^ Manually selected pathways with significant enrichment per cluster were plotted using Scanpy’s matrixplot().

### Neighborhood Enrichment

For each tissue section, low-abundance cell types (those making up < 1% of the tissue section) were removed. Squidpy was used to calculate spatial neighbors with Delaunay triangulation, and the squidpy.gr.nhood_enrichment() function was used to compute neighborhood enrichment with 1000 permutations.^116^ The Z-score matrix per section was averaged across sections from each patient, and the resulting patient-level Z-score matrices were then combined using Stouffer’s method (sum of the Z-scores divided by the square root of the number of patients). The resulting Z-score matrix was plotted using squidpy.pl.nhood_enrichment(), with NaN values set to zero.

### Moran’s I Spatial Autocorrelation

The Moran’s I score was calculated for each gene of the G4X breast panel in each tissue section using squidpy.^116^ Squidpy was used to calculate spatial neighbors with Delaunay triangulation. Next the squidpy.gr.spatial_autocorr() function was used with mode = “moran” and 100 permutations. The per-gene results were clustered using Seaborn’s clustermap() function using the Ward method on Euclidean distance.^117^ Significance was not able to be plotted using this method, but full results with p-values are shown in **Supplementary Table 7**.

### Immunohistochemistry

FFPE blocks from matching patients were obtained from our Institution’s Pathology Department. IHC was performed as previously described.^118^ Briefly, sections were deparaffinized in xylene, rehydrated in ethanol, and washed. Basic (DES, THSD4, NPM1), or citrate (c-JUN) unmasking was performed in pressure cooker for 30 minutes. After cooling, slides were washed in TBST and incubated in 3% hydrogen peroxide and then blocked. Slides were then incubated at 4°C overnight in the respective primary antibody (DES Monoclonal Antibody, Invitrogen Cat #MA5-13259; THSD4 Rabbit PolyAb, Proteintech Cat #20619-1-AP; c-JUN Rabbit mAb, Cell Signaling Technology Cat #9165S; NPM1 Monoclonal Antibody, Invitrogen Cat #32-5200). Slides were then washed, incubated in HRP-conjugated secondary antibody for 30 mins, washed and developed using DAB solution and mounted.

For NPM1 quantification, necrotic regions were manually identified in QuPath v0.6.0.^119^ Two concentric peri-necrotic regions were generated extending 0–50 µm (inner region) and 50–100 µm (outer region) from the necrotic boundary. Nuclear-based positive cell detection was performed using the Nucleus DAB Optical Density Mean measurement for NPM1 staining. A positive threshold cutoff of 1.75 was manually selected and applied across all biological replicates (n = 4). The percentage of positive cells within each region was quantified and compared between paired inner and outer regions. Statistical analysis was done using a paired t-test. Statistical analysis and bar plot generation were performed in GraphPad Prism 11.0.1.

### Bulk RNA Sequencing Survival Correlation

The Cancer Genome Atlas (TCGA) database was queried for human sarcoma bulk RNA sequencing datasets, and the expression and clinical metadata for each cohort were downloaded for further analysis. For the ULMS cohort (n=22), leiomyosarcoma (LMS) samples were filtered for tumors from the uterus, corpus uteri, or myometrium. For the LMS cohort (n=83), all cases labeled as leiomyosarcoma were included. For the sarcoma cohort (n=301), all sarcoma samples available through the TCGA were included, namely angiosarcoma, chordoma, clear cell sarcoma of the kidney, dedifferentiated liposarcoma, fibromyxosarcoma, leiomyosarcoma, myxoid leiomyosarcoma, osteosarcoma, synovial sarcoma, spindle cell sarcoma, undifferentiated sarcoma, gastrointestinal stromal tumor, malignant fibrous histiocytoma, malignant peripheral nerve sheath tumor, alveolar rhabdomyosarcoma, chondrosarcoma, epithelioid sarcoma, and well-differentiated liposarcoma.

For each tumor cell state identified on scRNA-seq, the top 50 differentially expressed genes with the highest rank_genes_groups() scores that were also present in the bulk expression data and had significant adjusted p-values (p_adj_ < 0.05) were compiled as the representative signature for that cluster. Using the gseapy package,^111^ Single-sample gene set enrichment analysis (ssGSEA) was then performed for each bulk RNA-seq sample using these signatures. This analysis generated a normalized enrichment score (NES) for each cluster within each sample. Bulk samples were then ranked by NES. Samples with NES values in the top and bottom quartiles for each state were used for Kaplan-Meier survival curves and log-rank testing.

### Drug Response Prediction Analysis

Cancer cell line drug response data used for prediction analyses were obtained from the Broad DepMap repository.^120^ Drug screening data were originally generated as part of the Cancer Therapeutics Response Portal version 2 (CTRPv2).^121^ We used the Simplicity pipeline–processed dataset to obtain normalized area-under-the-curve (nAUC) values.^122^ Cancer cell line transcriptomic data were originally generated through the Cancer Cell Line Encyclopedia (CCLE),^123^ and transcript-per-million (TPM)–normalized counts were downloaded from DepMap.

Drug responses for each tumor cluster were computed using the scIDUC (single-cell integration and drug response prediction) algorithm, as previously described.^87^ Briefly, a ranked list of drug response-relevant genes (DRGs) was generated for each drug using screening data from CTRPv2 (normalized area under the dose–response curve, nAUC). Both bulk RNA-sequencing (cancer cell lines, CCLs) and scRNA-seq datasets were restricted to the same DRGs to focus on drug-informative signals. Bulk and single-cell transcriptomes were integrated into a shared low-dimensional space using canonical correlation analysis, which preserved common expression patterns while reducing noise. Regression models (kernel regression) were then trained on bulk embeddings with CCL drug response values as outcomes, and the resulting coefficients were applied to single-cell embeddings to infer cell-level drug sensitivities. The drug response for a given cell cluster was defined as the mean nAUC of its cells. Subsequently, clusters were assigned a sensitivity rank from 1 to 8 for each drug, based on these mean values, where a rank of 1 indicates the highest sensitivity (lowest nAUC).

### Statistical Analysis

Statistical testing for all scRNA-seq analyses were based on default methods available for all functions described above. For TCGA-derived bulk RNA sequencing data, statistical testing was performed for the Kaplan-Meier survival plots using the log-rank test.

### Graphics and Illustrations

Graphical illustrations were created with BioRender. Adobe Illustrator 2025 and PowerPoint were used for figure layouts and formatting.

## Data Availability

Raw scRNA-seq and spatial data were deposited on GEO, and all data will be made publicly available at the time of publication of this work. 3SEQ data for correlations were downloaded from GSE45510.

## Acknowledgements

First and foremost, we would like to thank the many extraordinary and inspiring patients with uterine sarcoma who contributed to this study over the past several years. This study would not have been possible without them. We would like to dedicate this study to these patients and to others like them who have been diagnosed with rare diseases. This work was made possible by the contributions of Retro Biosciences team, especially the guidance of Jared Tumiel, J.N. Rashida Gnanaprakasam, Jacqueline A Larouche, Tobias Fehlmann, and Alexandre Trapp. We would also like to acknowledge the contributions of the Singular team, especially the guidance from Jordan Williams and Darius Fugere. D.D. was supported by funding from the Damon Runyon Cancer Research Foundation, the American Cancer Society, the National Institutes of Health (NIH) Loan Repayment Program, and the John and Marva Warnock Faculty Scholar Award. M.K. was supported by the Division of General Surgery, Department of Surgery, School of Medicine, Stanford University and the Advanced Residency Training at Stanford Program. M.K. and J.P.A. were supported by the Transplant Tissue Engineering Center of Excellence at Stanford University. This work was supported by the Stanford Functional Genomics Facility and Stanford Cancer Institute. Computing for this project was performed on the Stanford SCG Bioinformatics Cluster (RRID:SCR_026876), owned by the Stanford Genomics Bioinformatics Service Center (RRID:SCR_023340) and operated by Stanford Research Computing (RRID:SCR_023413). We would like to acknowledge Chris Jeon (ORCID: 0009-0007-4068-7598) of Stanford Research Computing (RRID:SCR_023413), who provided assistance with setup and troubleshooting of the GPU cluster.

## Author contributions

D.D. conceived and supervised the project. M.K. and J.P.A designed and conducted most experiments, analyzed the data, and wrote the manuscript. B.R., L.W., and B.L. helped analyze the data. R.A.B.F., and A.G. performed experiments. W.B. and K.A.Y. contributed to initial manuscript and figure preparation. R.R.T., C.G., R.G., A.E.D., P.Y.X., A.T., B.S., and S.K.D. assisted with experimental and technical setup. B.L., O.D., N.Q.B., M.P., E.J.M., A.K., A.R.K., G.A.P., J.A.N., A.M.N., G.C., K.N.G., D.S.F., R.S.H., B.L., and M.T.L., provided conceptual inputs for the paper and contributed to manuscript revision. All authors read and approved the final manuscript.

## Extended data figure legends

**Extended Data Fig. 1.**
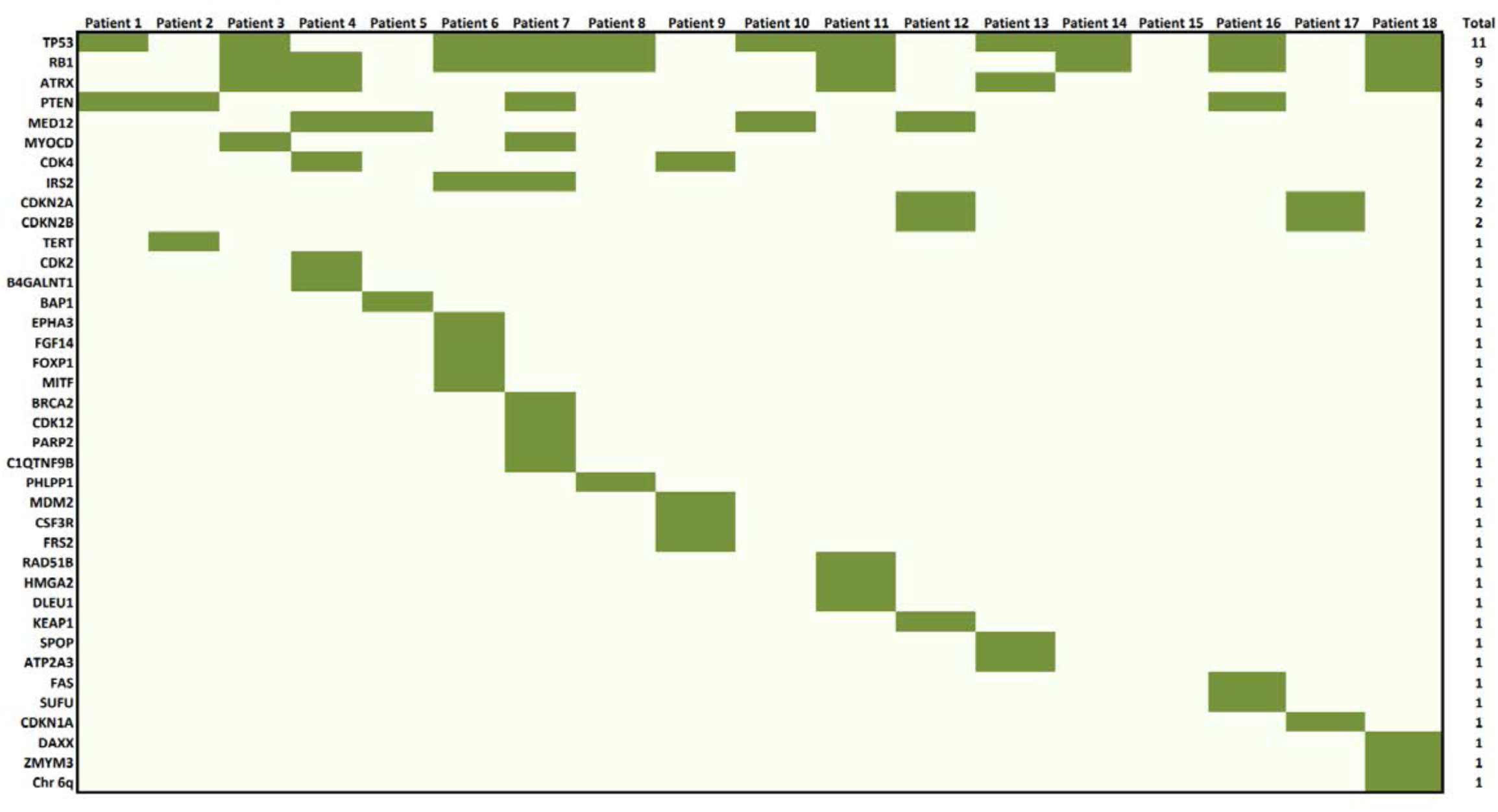
List of all mutations of known significance present across each patient.

**Extended Data Fig. 2.**
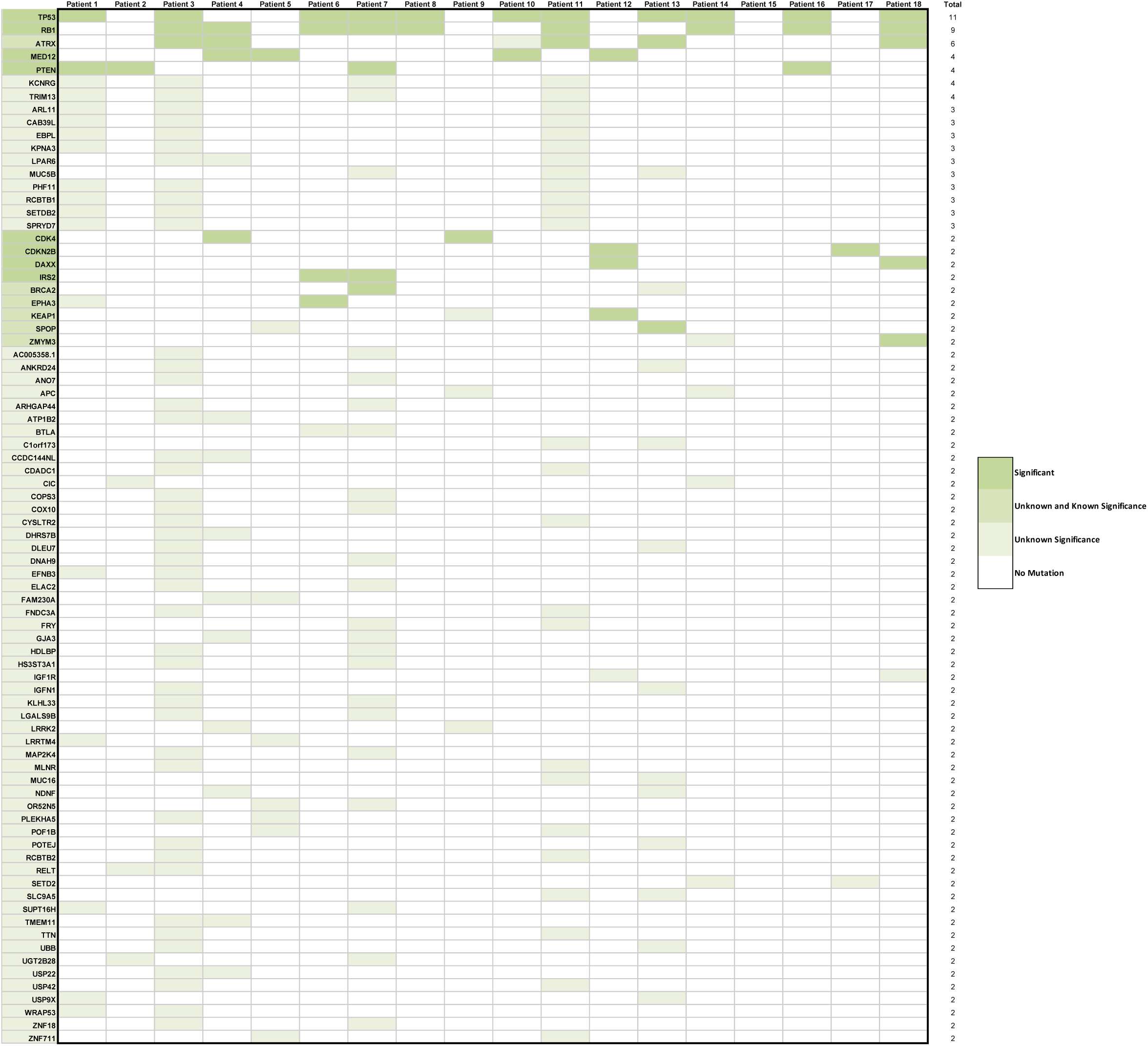
List of all variants of known and unknown significance present in at least two or more patients, colored by whether the variants were of known or unknown significance.

**Extended Data Fig. 3.**
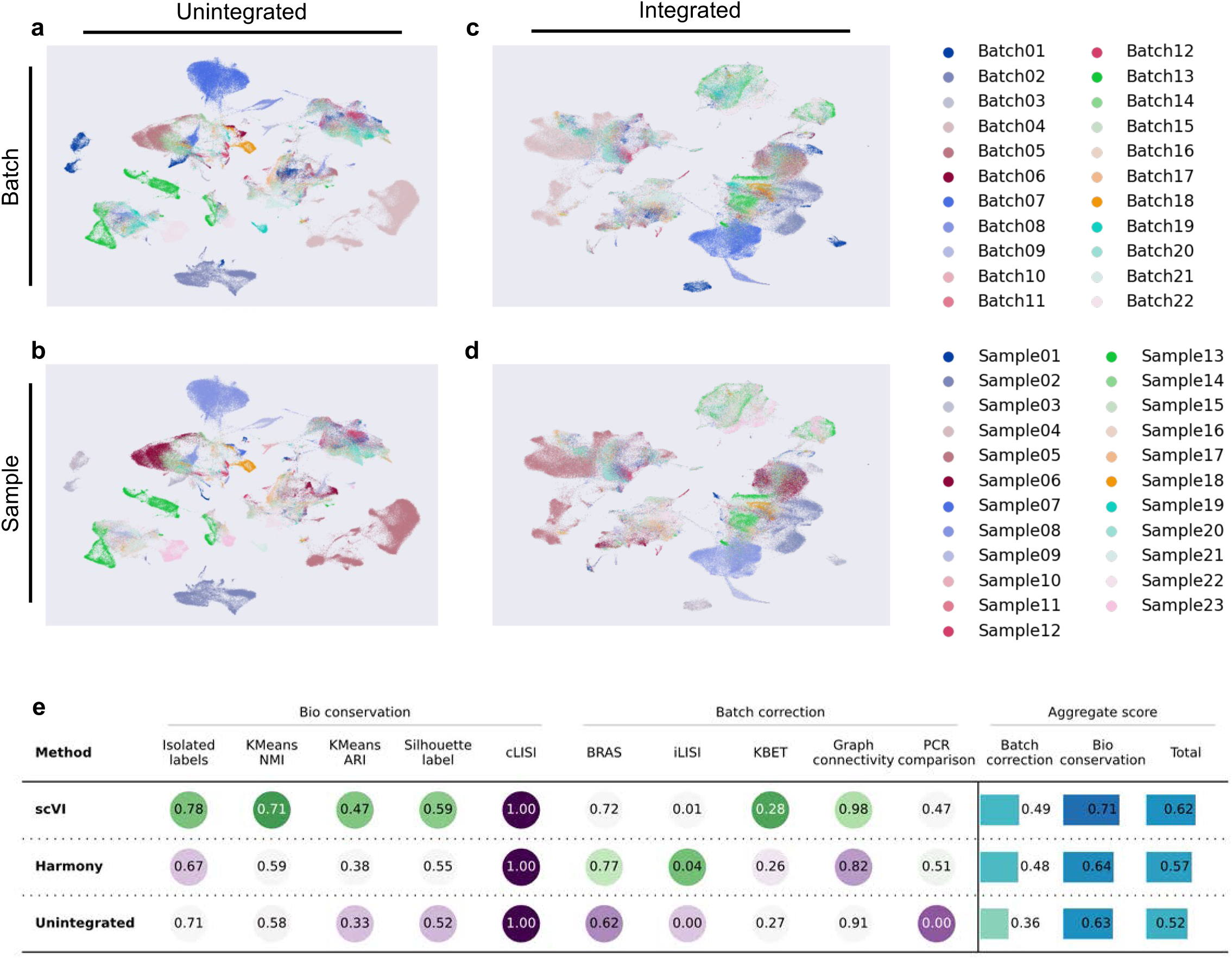
**a**, Unintegrated UMAP of all cells, colored by batch. **b**, Unintegrated UMAP of all cells, colored by sample. **c**, Integrated UMAP of all cells, colored by batch. **d**, Integrated UMAP of all cells, colored by sample. **e**, Results of scIB demonstrating biological conservation, batch correction, and aggregate score across integration methods.

**Extended Data Fig. 4.**
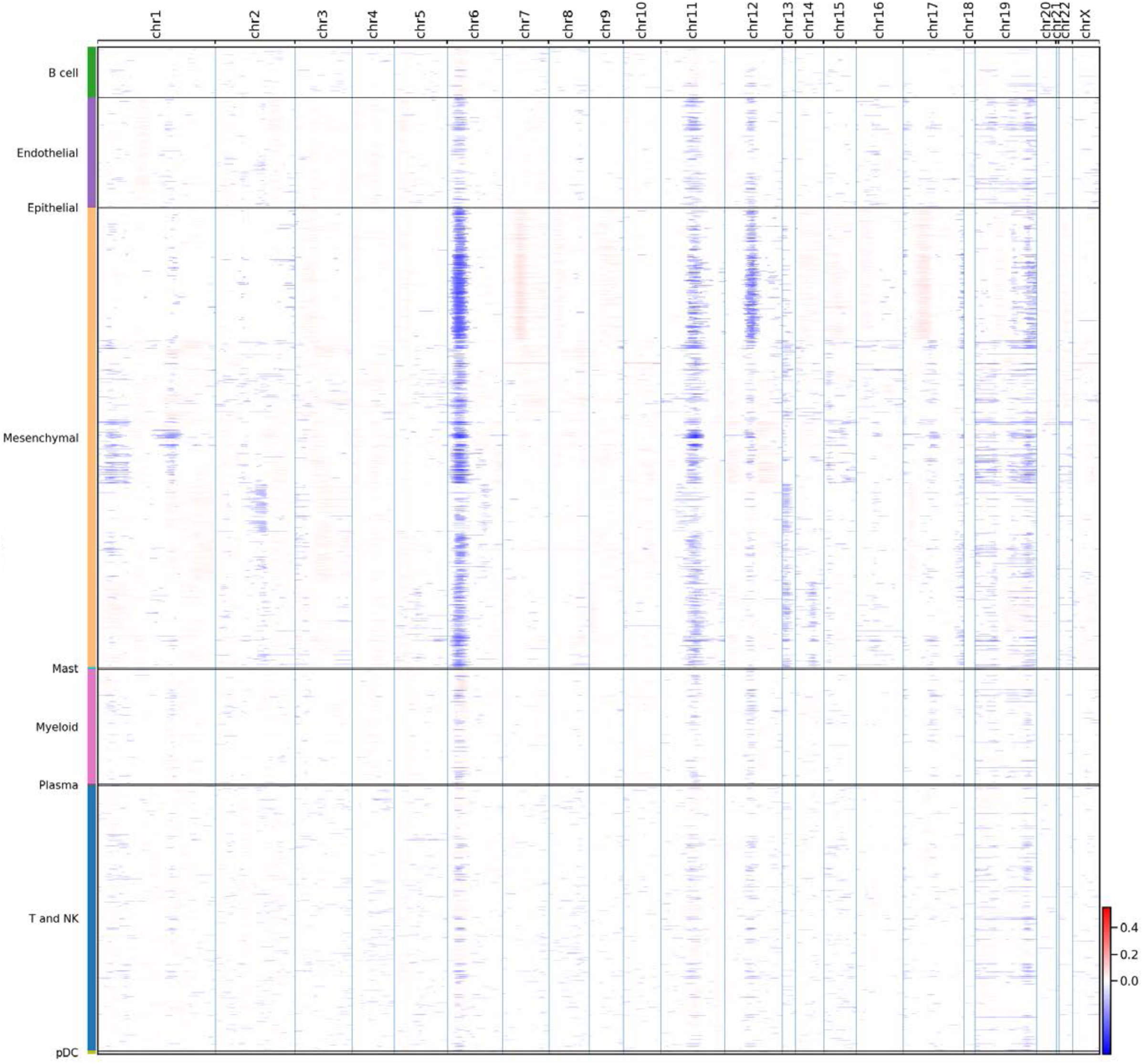
InferCNV results displayed as a heatmap across each chromosome for all cells.

**Extended Data Fig. 5.**
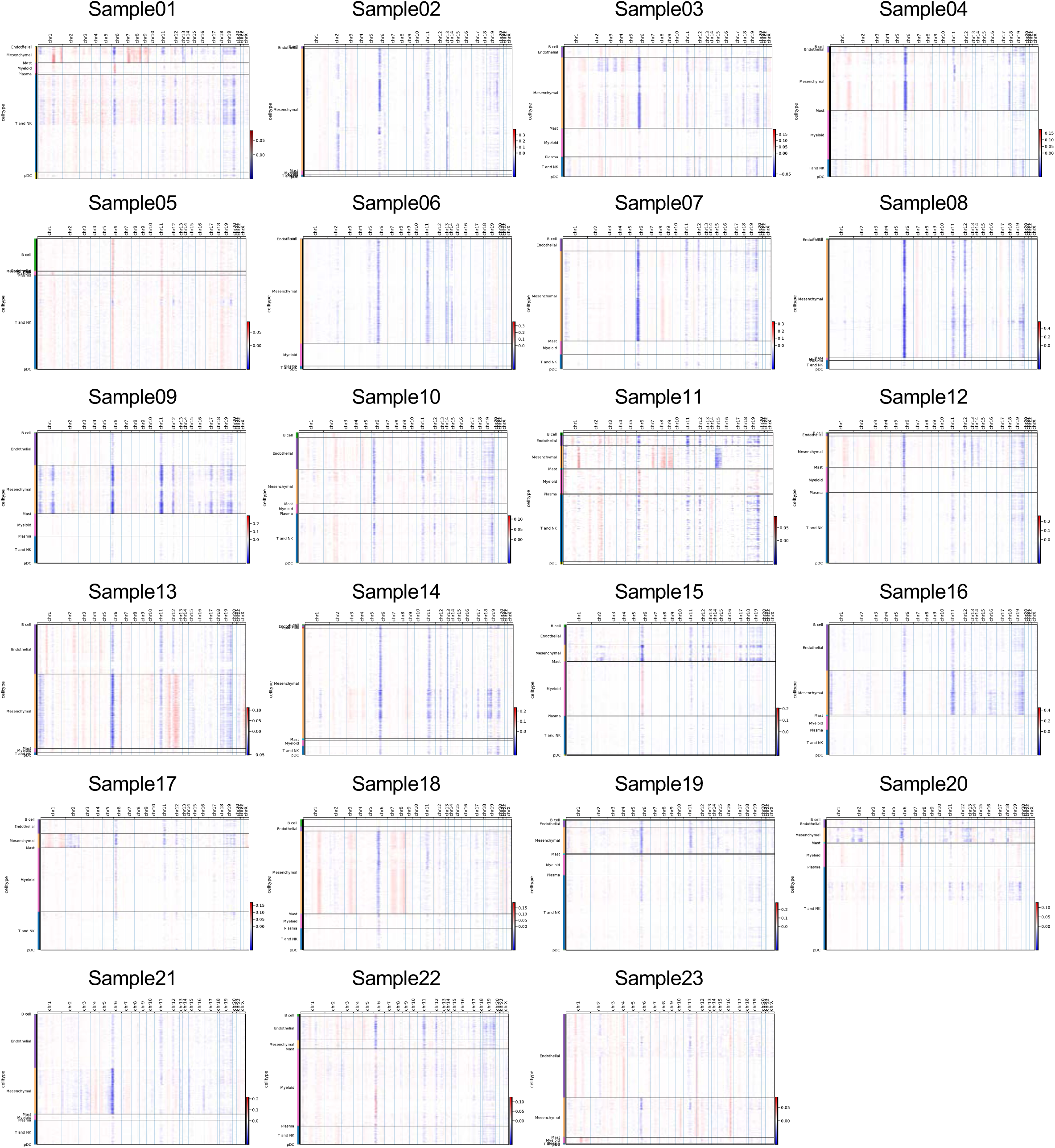
InferCNV results displayed as heatmaps per sample, across each chromosome for all cells.

**Extended Data Fig. 6.**
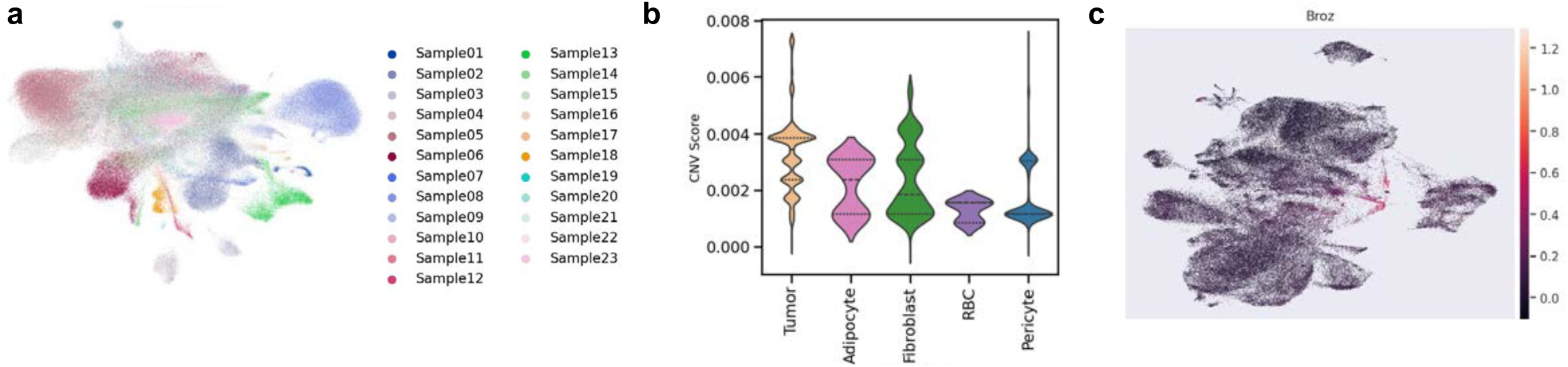
**a**, UMAP of all cells clustered by CNV, colored by sample. **b**, Violin plot of CNV scores across cell types in the mesenchymal subset. **c**, Integrated UMAP of the mesenchymal subset cells, colored by a composite of genes previously shown to distinguish fibroblasts from sarcoma tumor cells by Broz, et al. *Nat Comm*, 2024.

**Extended Data Fig. 7.**
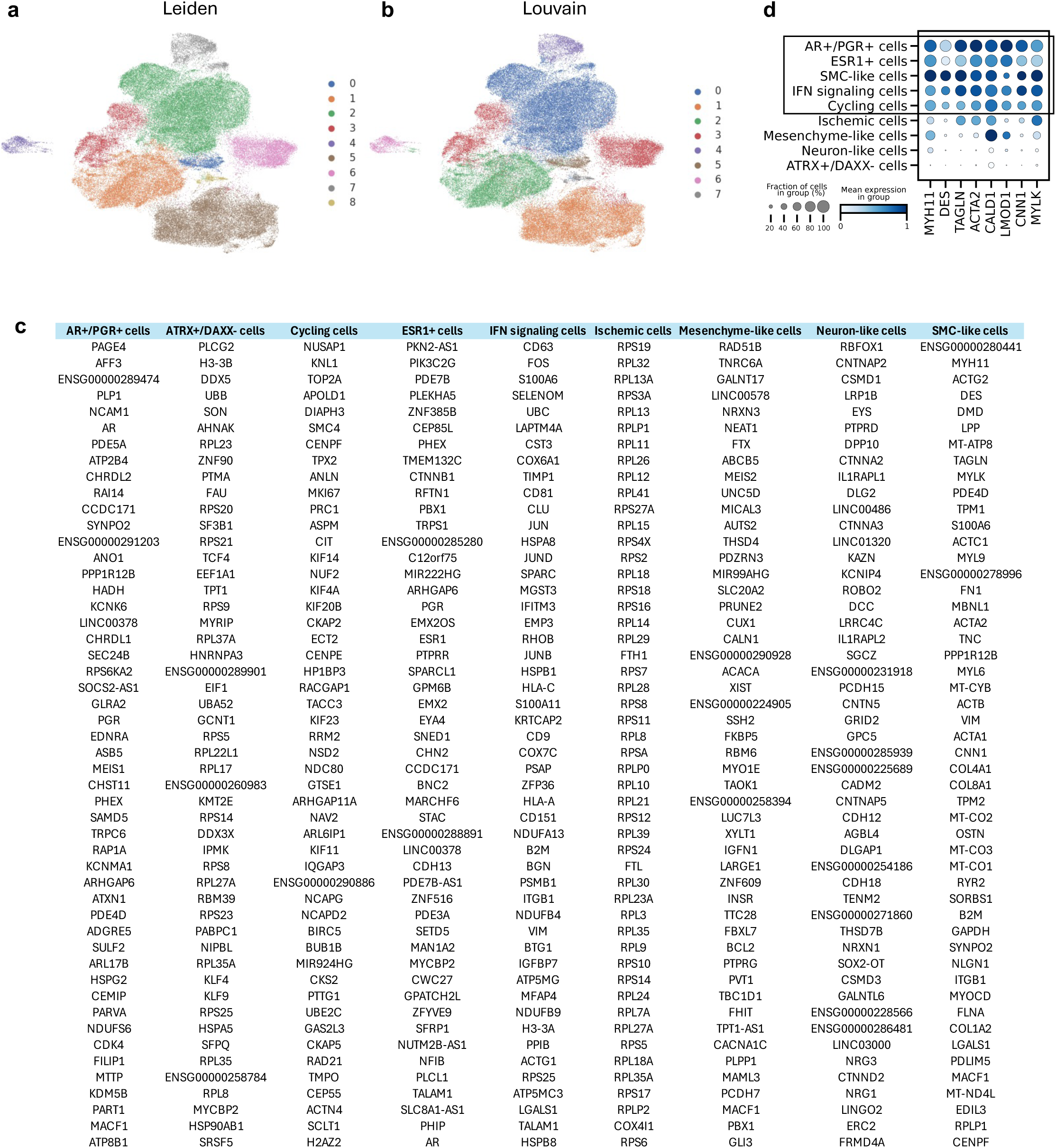
**a**, Integrated UMAP of tumor cell subset clustered by Leiden clustering, colored by cluster. **b**, Integrated UMAP of tumor cell subset clustered by Louvain clustering, colored by cluster. **c**, List of top 50 differentially expressed scanpy genes per tumor cell state. **d**, Dot plot of tumor cell states showing expression of various canonical SMC markers.

**Extended Data Fig. 8.**
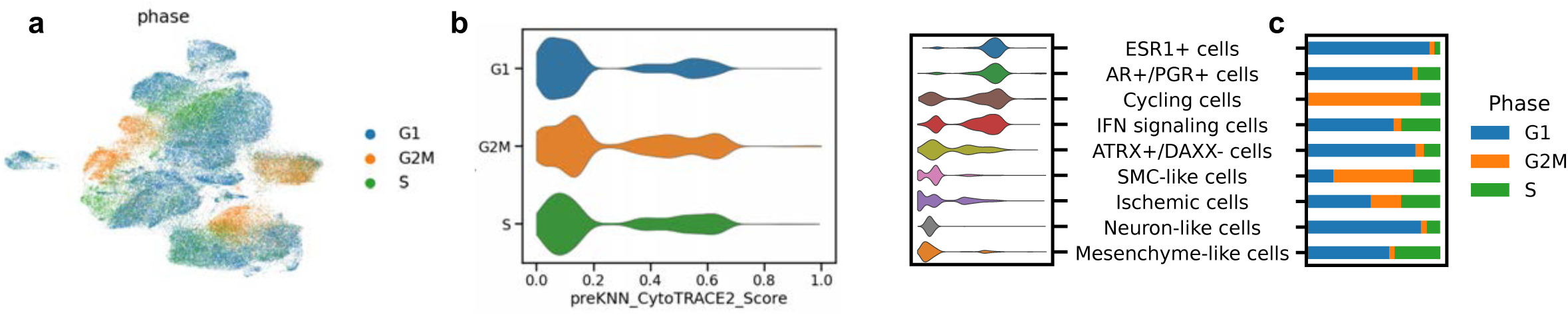
**a**, Integrated UMAP of tumor cell subset colored by calculated cell cycle phase. **b**, Violin plot depicting CytoTRACE 2 pre-KNN scores by cell cycle phase. **c**, Bar plot depicting proportion of cells within each tumor state predicted to be in each cell cycle phase, with the violin plot from Fig 2d showing CytoTRACE 2 pre-KNN scores on the left for reference.

**Extended Data Fig. 9.**
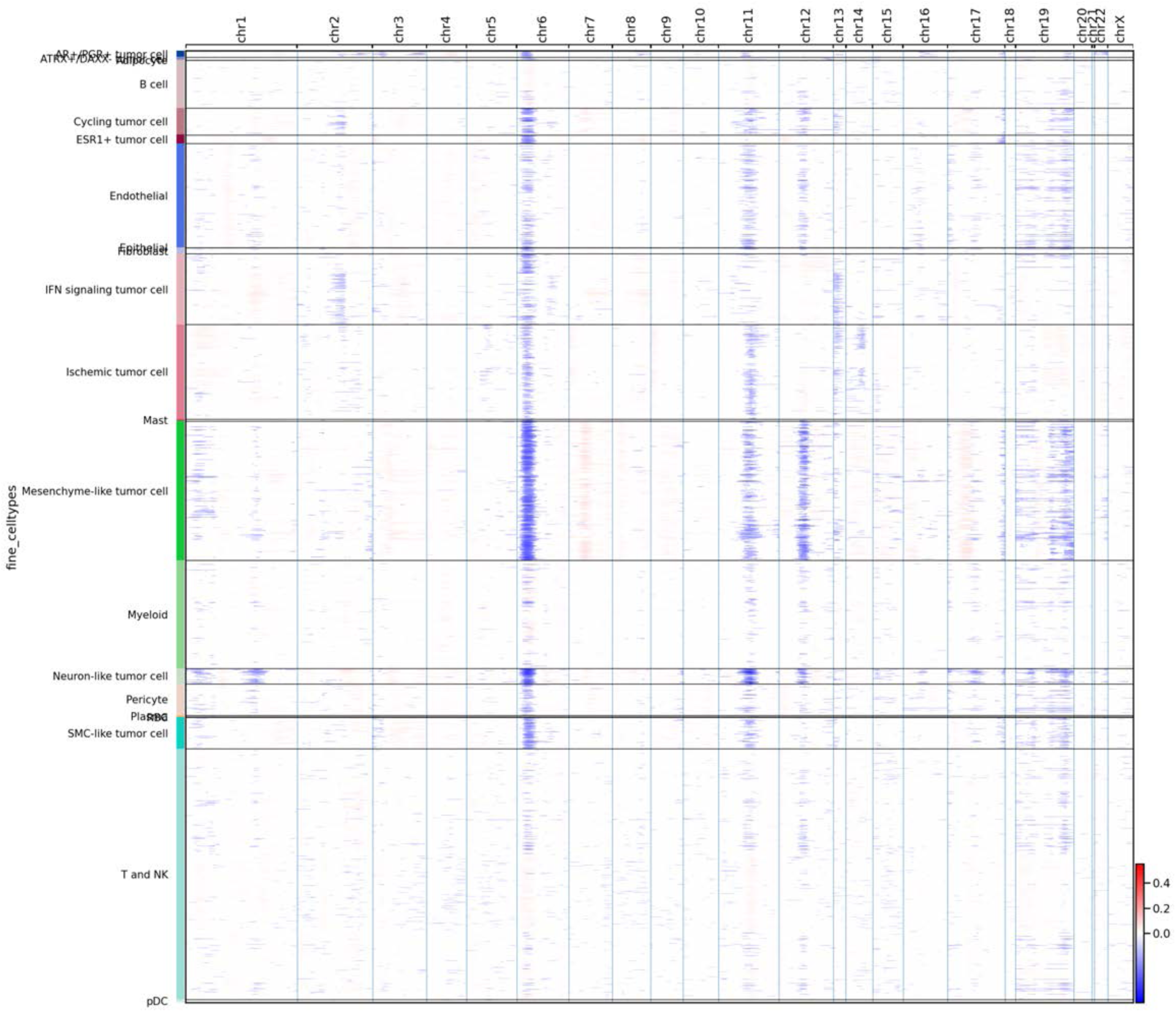
InferCNV results displayed as a heatmap across each chromosome for all cells with tumor cell states annotated.

**Extended Data Fig. 10.**
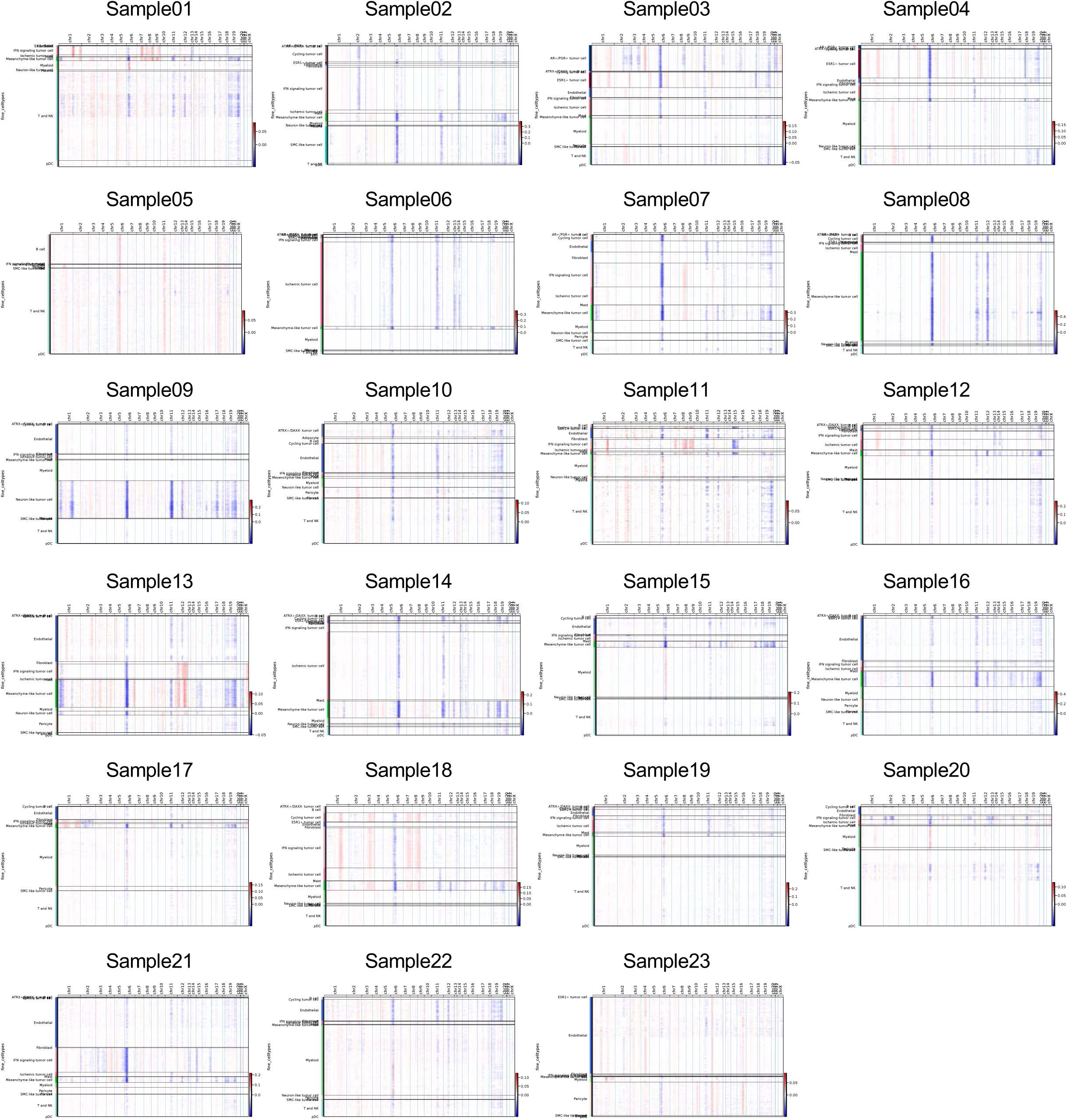
InferCNV results displayed as heatmaps per sample, across each chromosome for all cells, with tumor cell states annotated.

**Extended Data Fig. 11.**
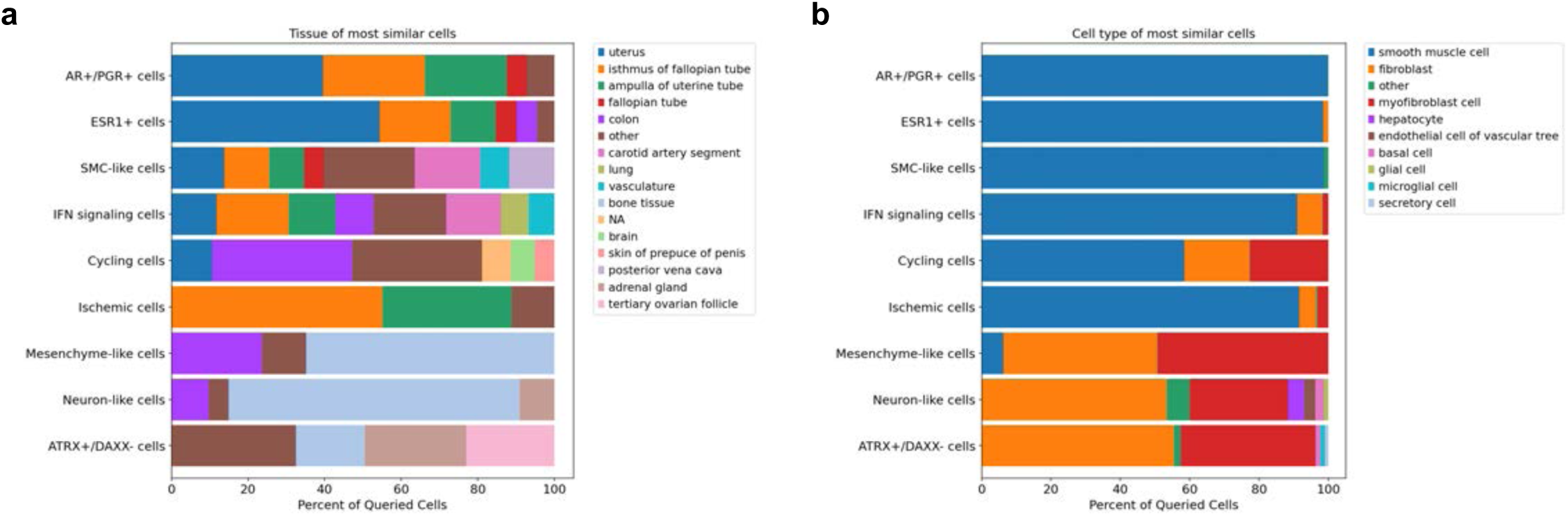
Bar plots representing results from scimilarity cell search, listing cell type **(a)** and tissue **(b)** of the top 10,000 cells that are most similar to each tumor cell state.

**Extended Data Fig. 12.**
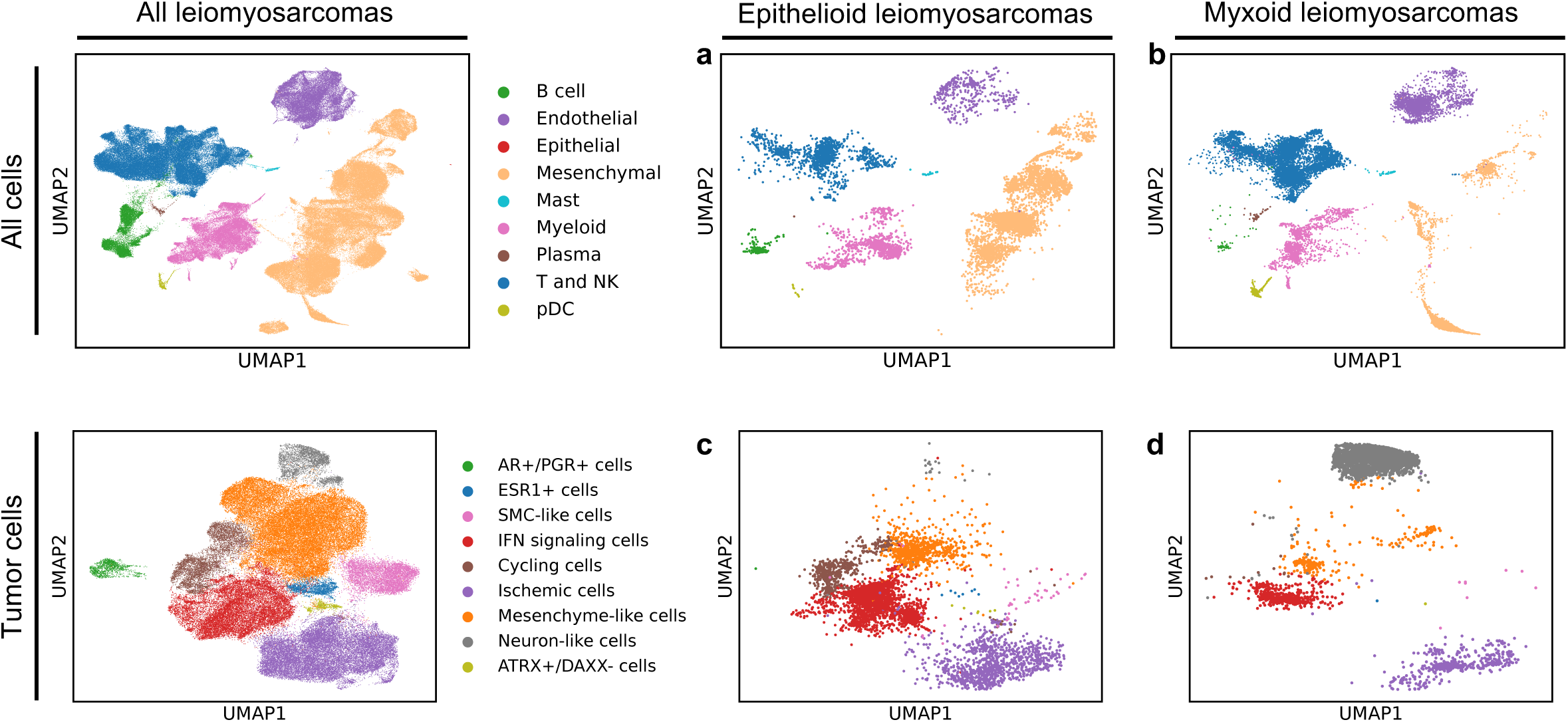
**a**, Integrated UMAP of all cells, showing only cells from epithelioid leiomyosarcoma samples, colored by cell type. **b**, Integrated UMAP of all cells, showing only cells from myxoid leiomyosarcoma samples, colored by cell type. **c**, Integrated UMAP of tumor cells, showing only cells from epithelioid leiomyosarcoma samples, colored by tumor cell states. **d**, Integrated UMAP of tumor cells, showing only cells from myxoid leiomyosarcoma samples, colored by tumor cell states.

**Extended Data Fig. 13.**
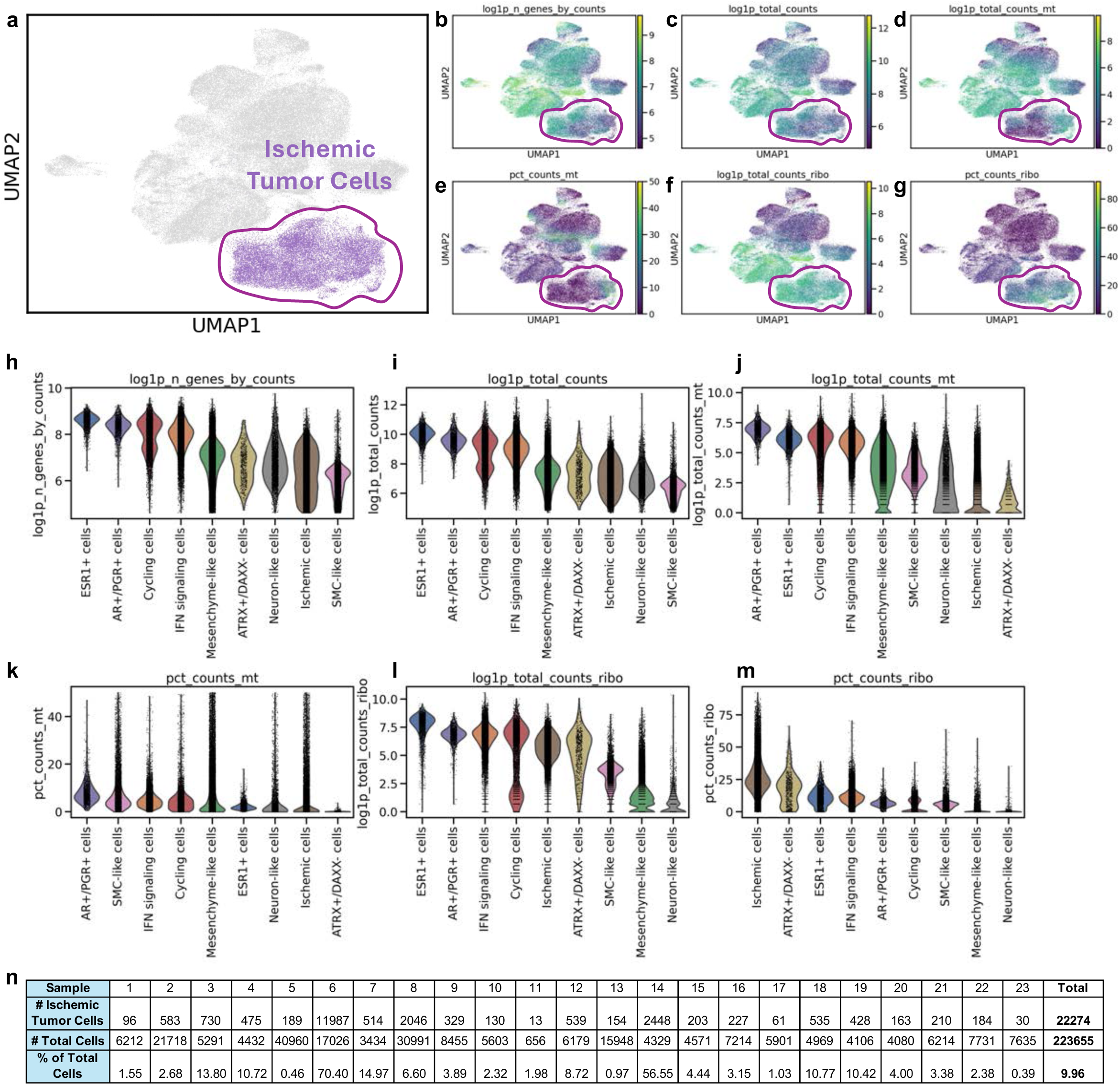
**a**, Integrated UMAP of tumor cell subset with Ischemic tumor cells highlighted. **b-g**, Integrated UMAPs of tumor cell subset colored by **(b)** log transformed number of genes per cell, **(c)** log transformed total counts, **(d)** log transformed mitochondrial gene counts, **(e)** percent mitochondrial counts, **(f)** log transformed ribosomal gene counts, and **(g)** percent ribosomal gene counts. **h-m**, Violin plots showing **(h)** log transformed number of genes per cell, **(i)** log transformed total counts, **(j)** log transformed mitochondrial gene counts, **(k)** percent mitochondrial counts, **(l)** log transformed ribosomal gene counts, and **(m)** percent ribosomal gene counts across tumor cell states. **n**, Table listing number of ischemic tumor cells, total number of cells, and percent of total number of cells comprised by ischemic tumor cells, per sample and the total across all samples.

**Extended Data Fig. 14.**
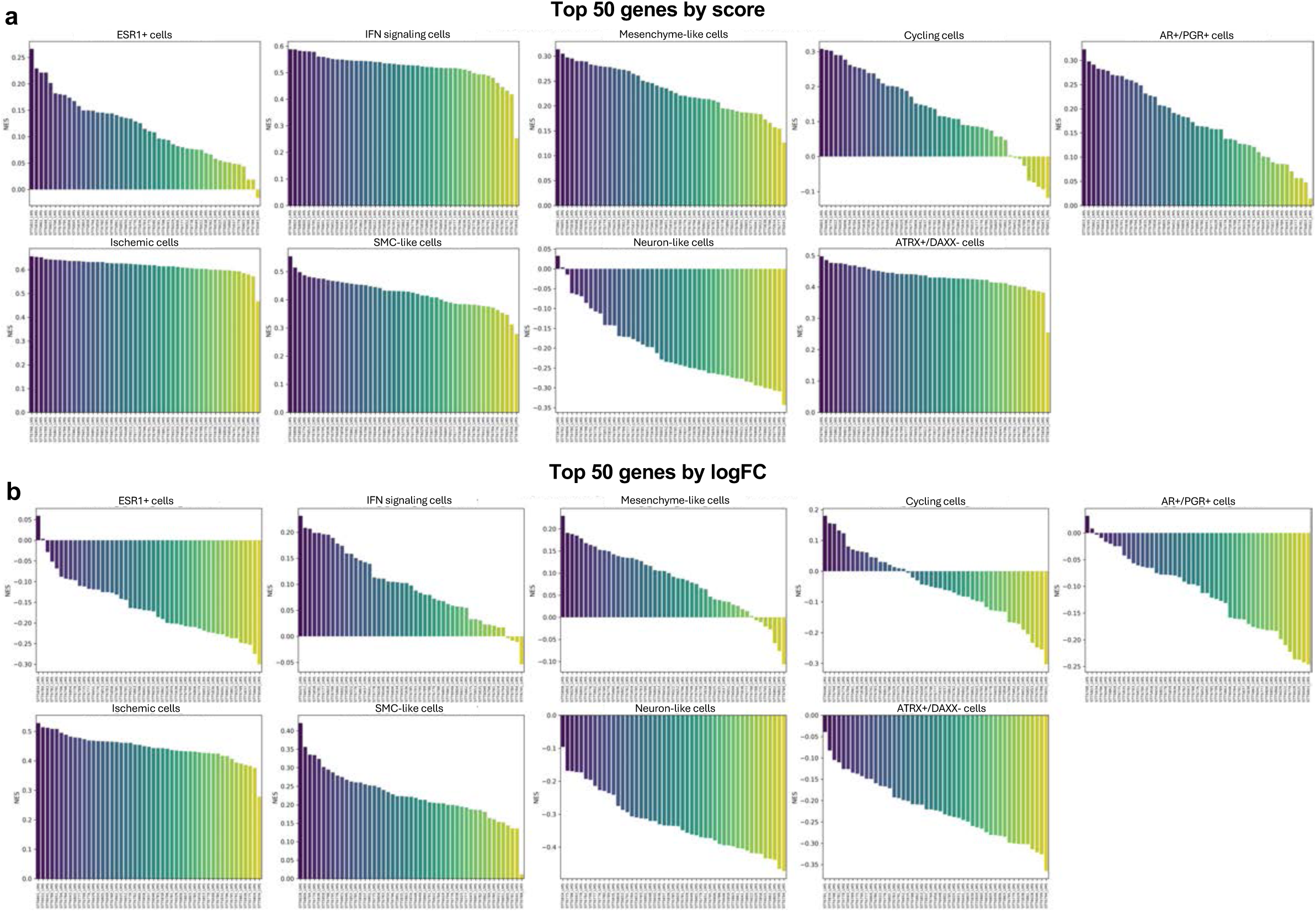
**a-b**, Bar plots displaying normalized enrichment scores from ssGSEA analysis of 3SEQ datasets of uterine leiomyosarcomas, from Guo et al. (*Clin Cancer Res*, 2015), with scores computed using top 50 genes with the highest scores **(a)** or highest log fold change **(b)**.

**Extended Data Fig. 15.**
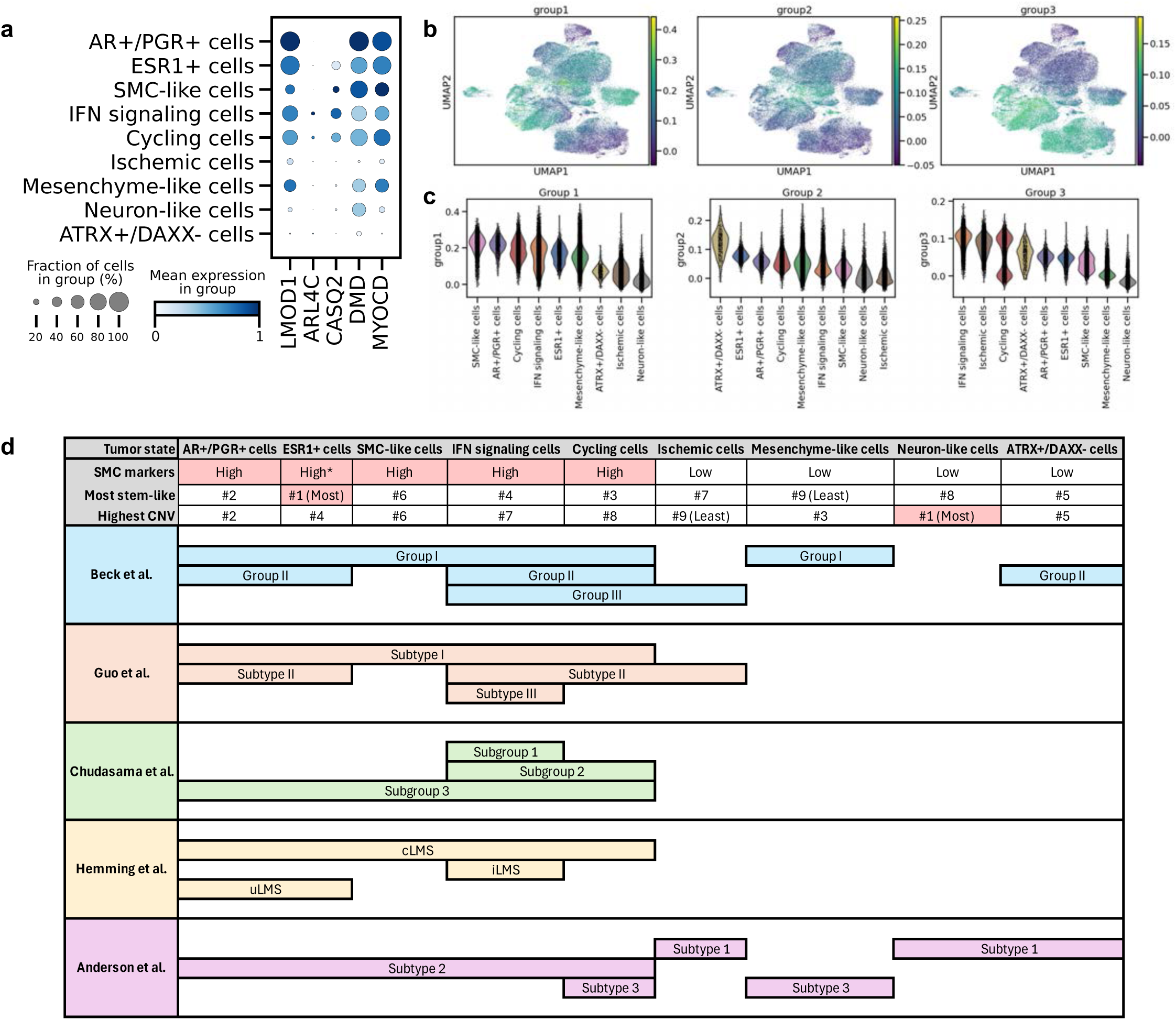
**a**, Dot plot of markers previously shown to be differentially expressed across leiomyosarcoma subtypes, plotted across tumor cell states. **b**, Integrated UMAPs of the tumor subset, colored by an aggregate score representing highly expressed genes in groups I-III described in Beck, et al. (*Oncogene*, 2009). **c,** Violin plots of the aggregate scores representing highly expressed genes in groups I-III described in Beck, at al. (*Oncogene*, 2009), plotted per tumor state. **d**, Diagram correlating previously described leiomyosarcoma subtypes with the ULMS tumor states described in this study

**Extended Data Fig. 16.**
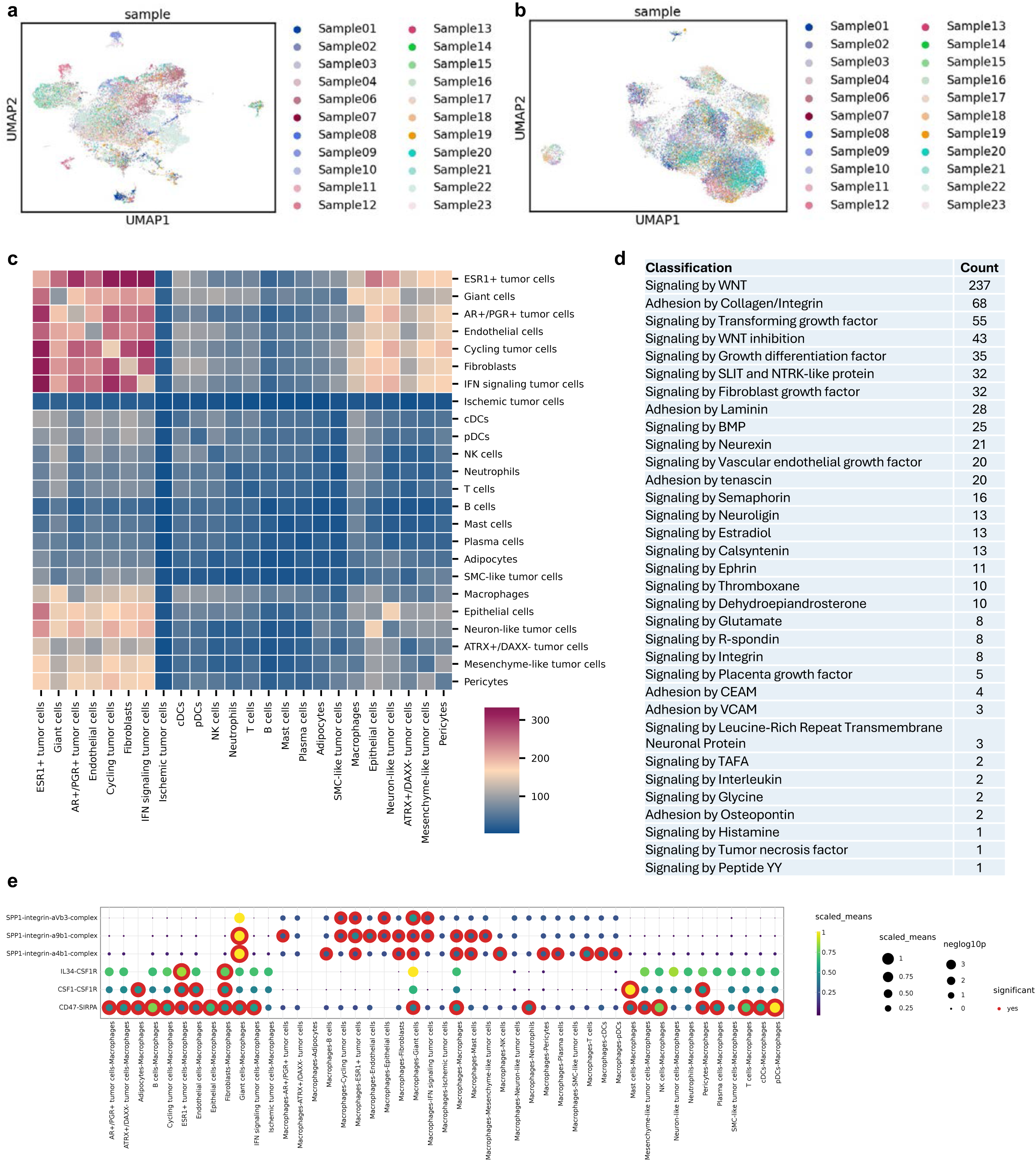
**a**, Integrated UMAP of myeloid cells colored by sample. **b**, Integrated UMAP of lymphoid cells colored by sample. **c**, Heatmap displaying sum of significant cell-cell interactions computed using CellphoneDB. **d**, Table showing the frequency of statistically significant pathway classifications with a minimum score threshold of 50, for cell-cell interactions between ESR1+ tumor cells and all other cell types, inferred using CellphoneDB. **e**, CellphoneDB dot plot showing selected macrophage interactions.

**Extended Data Fig. 17.**
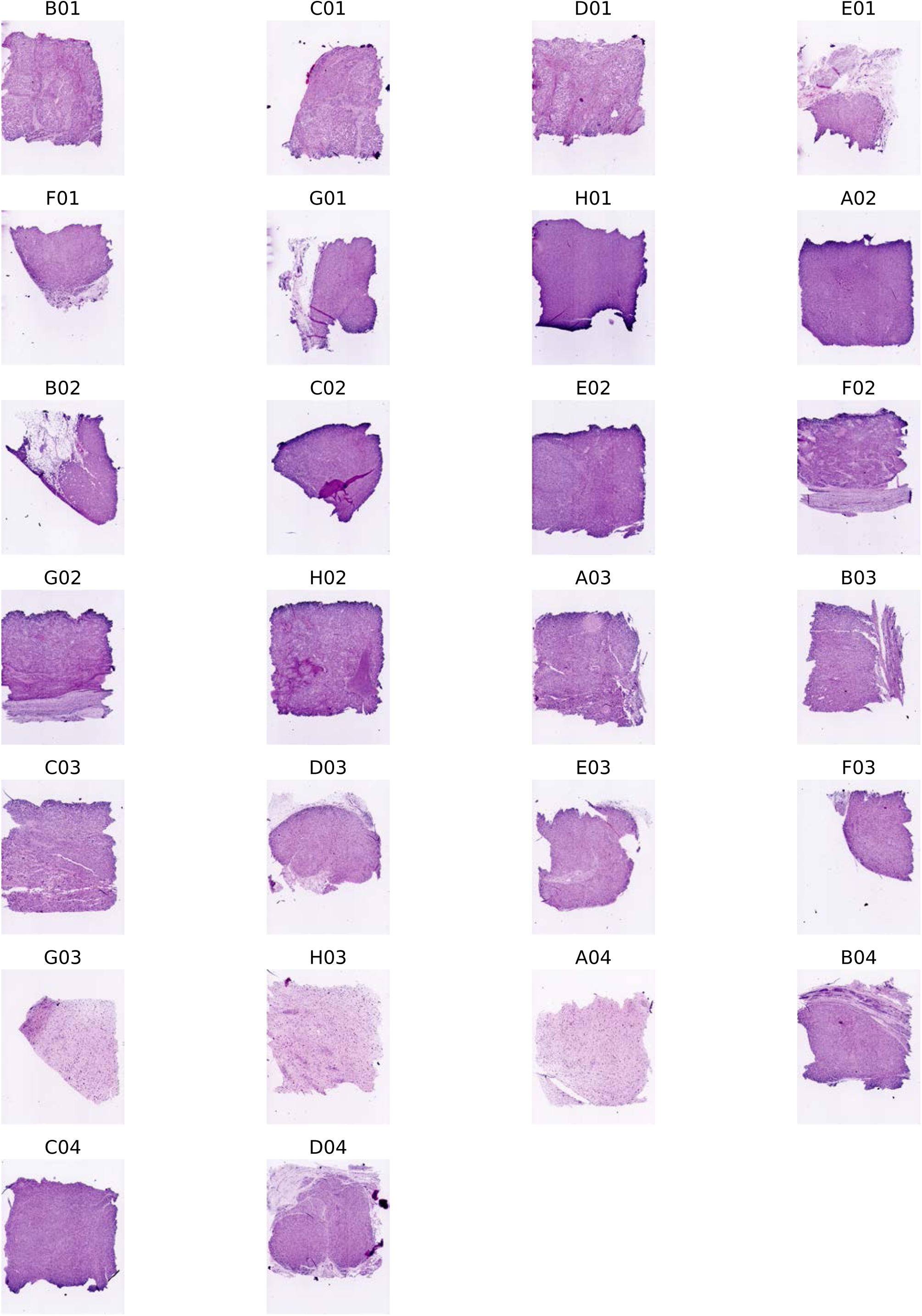
**a**, Fluorescent H&E of each section analyzed using the Singular G4X spatial platform.

**Extended Data Fig. 18.**
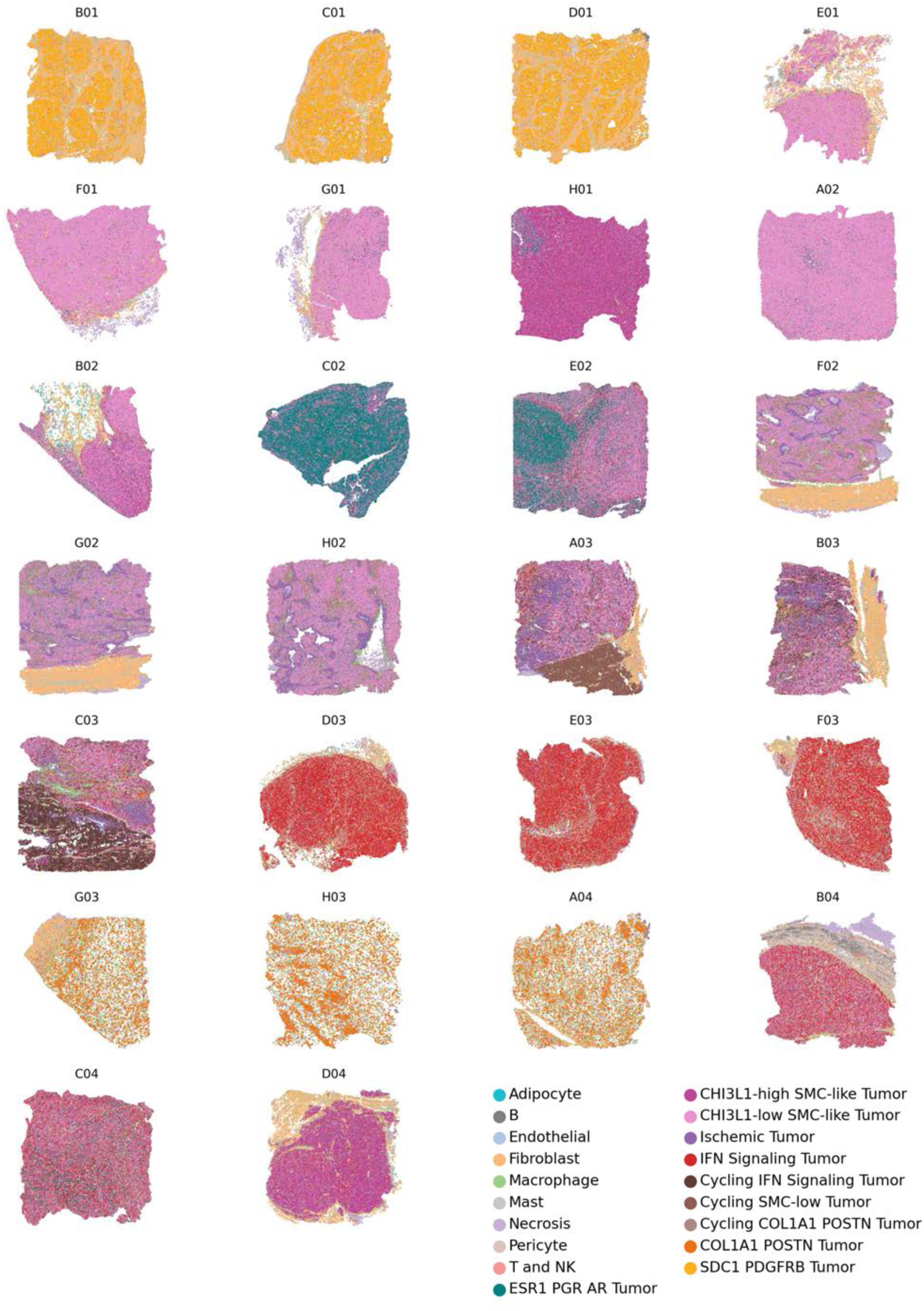
Cell type annotations presented spatially for each section analyzed using the Singular G4X spatial platform.

**Extended Data Fig. 19.**
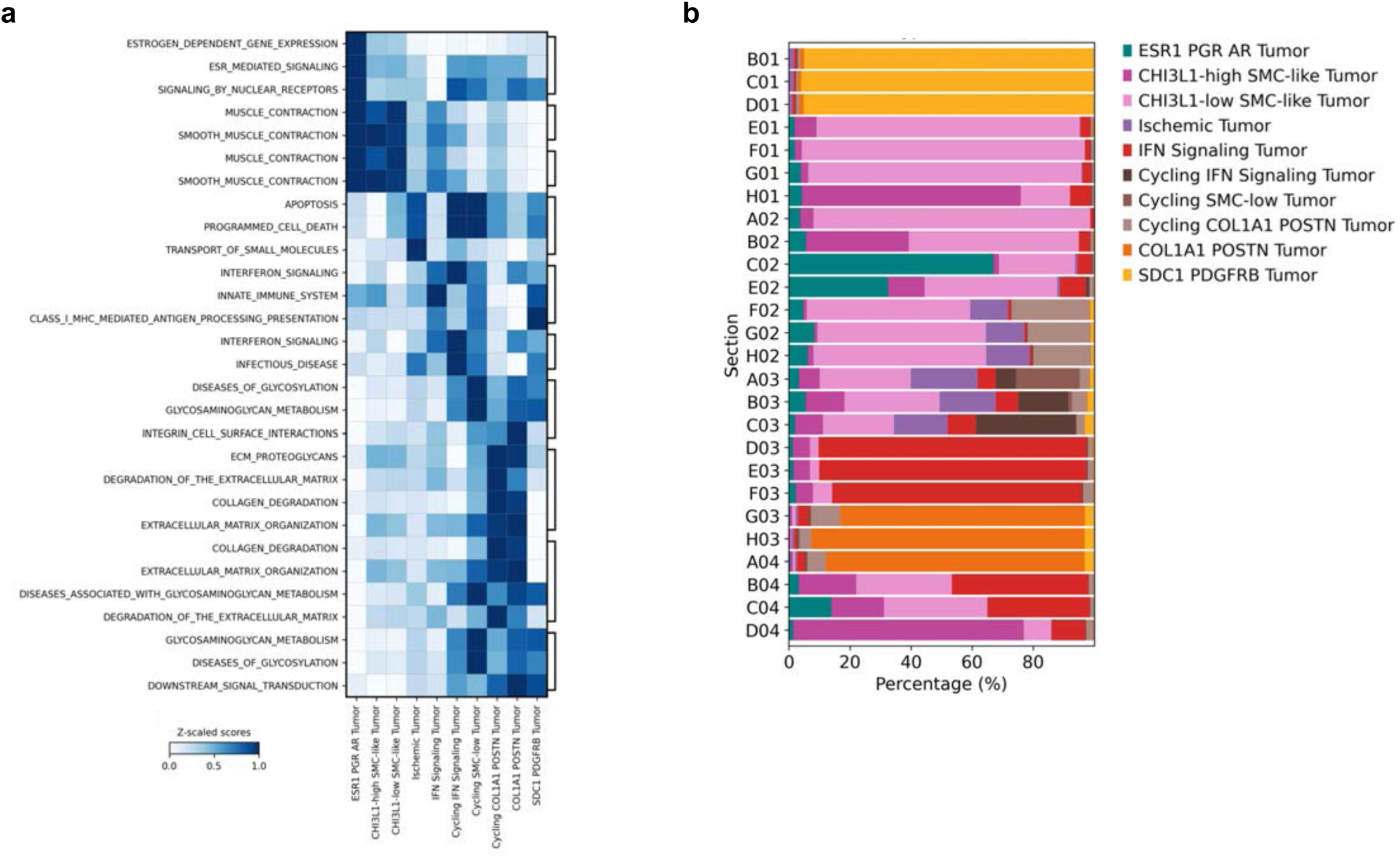
**a,** Matrix plot of selected reactome pathways enriched in each spatial tumor state. **b,** Bar plot displaying the proportion of each spatial tumor cell state present within each sample.

**Extended Data Fig. 20.**
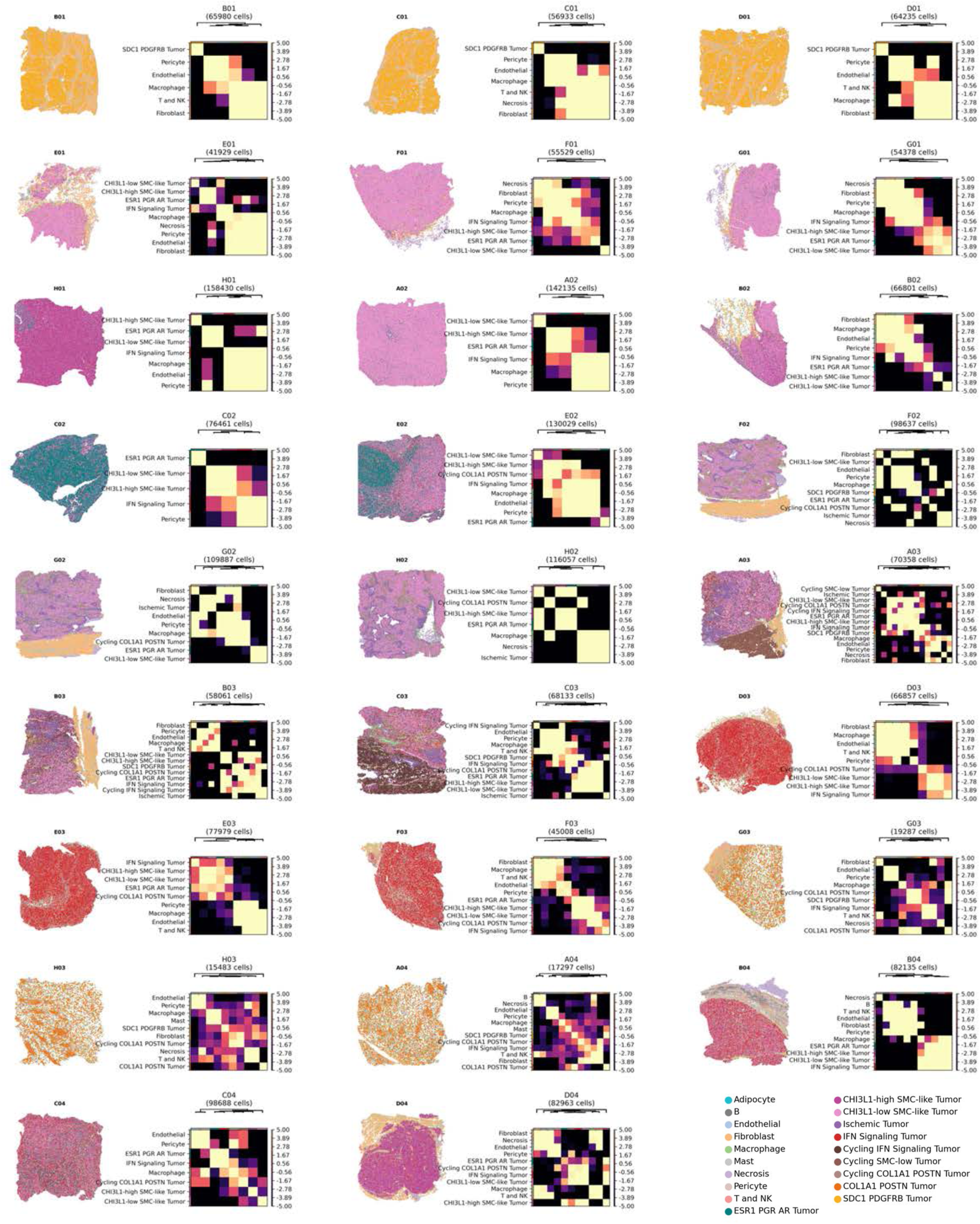
Matrix plot showing neighborhood enrichment analysis per sample for all tumor cell states, presented adjacent to each corresponding annotated section.

**Extended Data Fig. 21.**
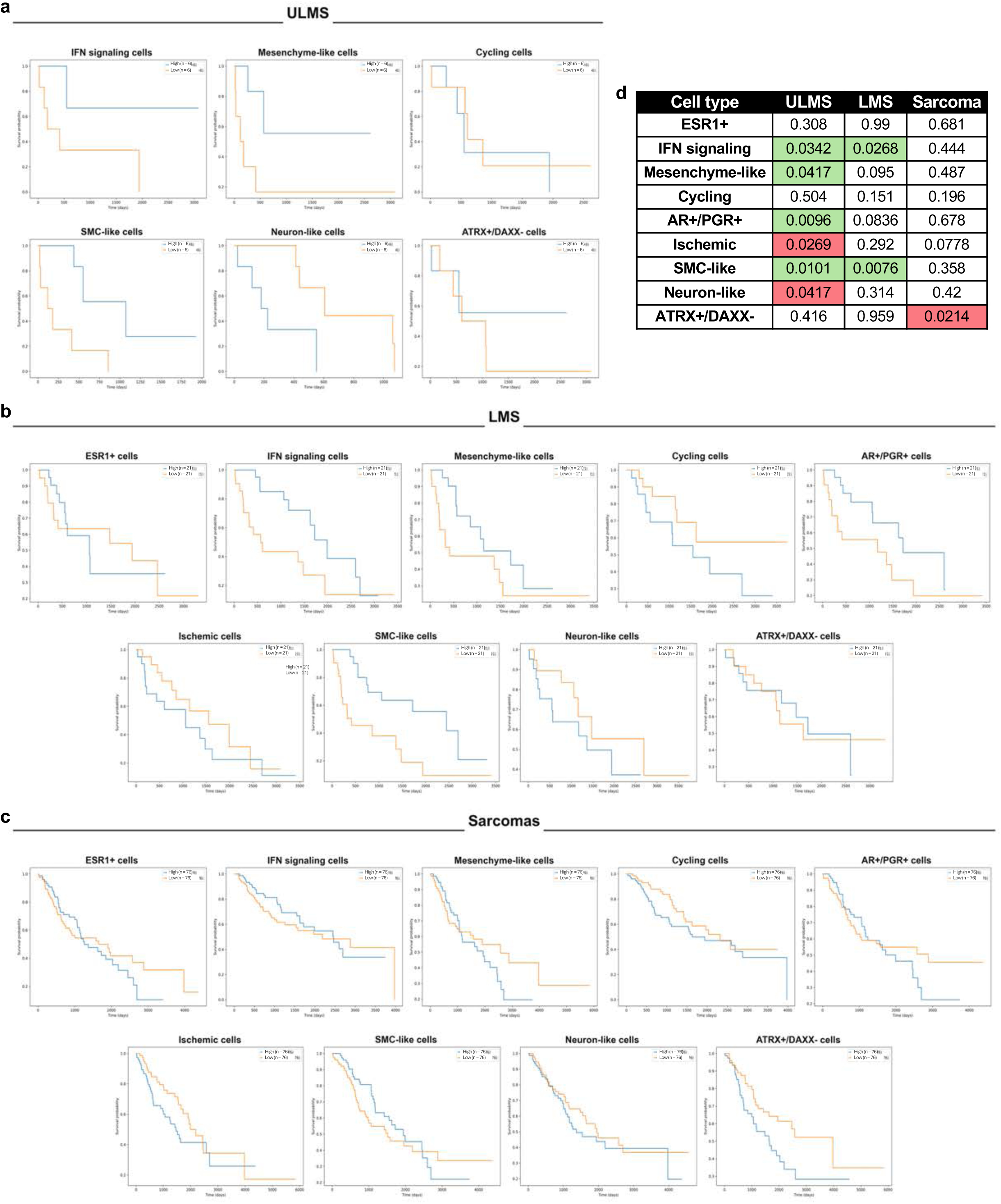
**a-c**, Kaplan-Meier curves of patients with high and low enrichment of individual tumor state signatures, using bulk RNA sequencing data from TCGA for **(a)** ULMS, **(b)** LMS, and **(c)** all sarcomas. **d**, P-values of Kaplan-Meier curves for each tumor cell state across ULMS, LMS, and all sarcomas from TCGA, colored by survival benefit (green), disadvantage (red), or non-significant value (grey).

**Extended Data Fig. 22.**
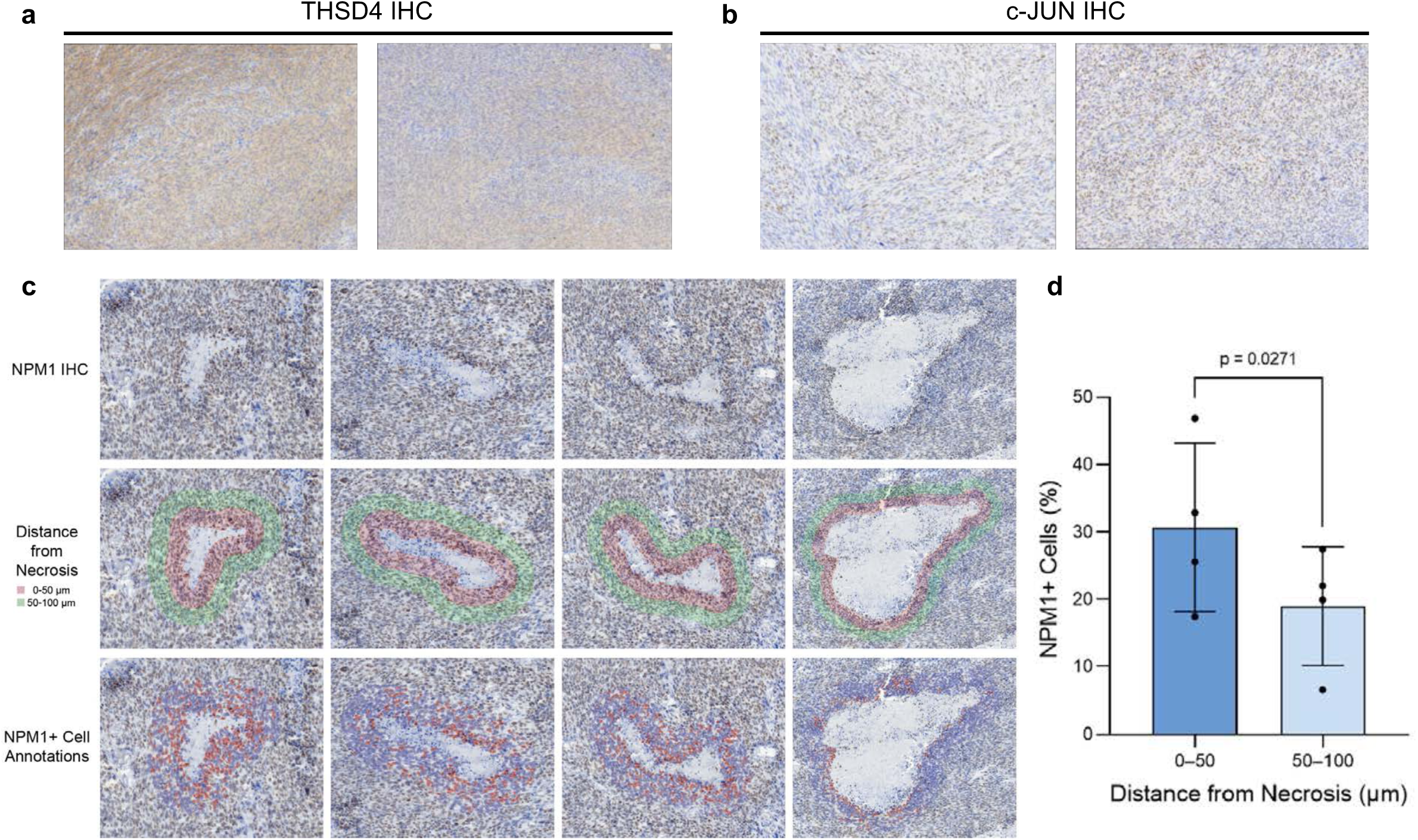
**a-b**, Representative IHC images from ULMS tumors from different patients staining positive for **(a)** THSD4 and **(b)** c-JUN. **c**, Representative images of necrotic areas with NPM1 IHC (top), with borders marking the areas quantified (middle), and cell detection outlines (bottom). **d,** Bar plot displaying a comparison of percentage of cells with positive NPM1 detection immediately adjacent to necrotic areas (0-50 μm from necrosis) and further away from necrotic areas (50-100 μm from necrosis). Individual points represent each necrotic area quantified (n=4), height of the bar represents mean, and error bars represent standard deviation.

**Extended Data Fig. 23.**
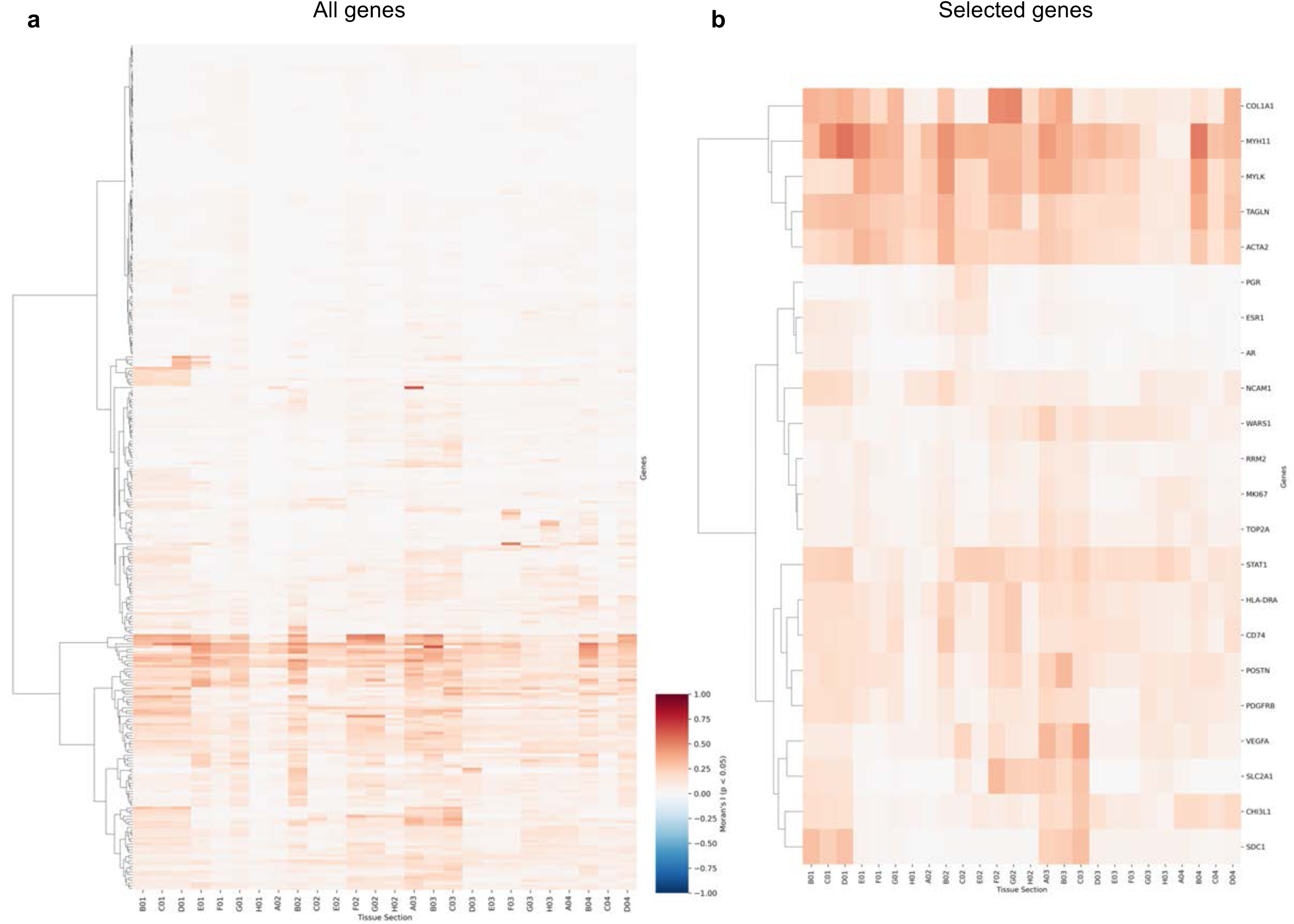
**a-b,** Clustered heat map depicting Moran’s I spatial autocorrelation results for **(a)** all genes, and **(b)** selected genes used to annotate each cluster. Each column is a spatial sample, and each row is a gene. Moran’s I of +1 (red) indicates clustering, -1 (blue) indicates dispersion, and 0 (white) indicates a random distribution.

## Supplementary table legends

**Supplementary Table 1.** Detailed characteristics of each patient included in this study.

**Supplementary Table 2.** Abbreviations of chemotherapy agents depicted in Fig. 1b.

**Supplementary Table 3.** List of variants of unknown significance derived from clinical mutation panel testing.

**Supplementary Table 4.** Scanpy differentially expressed genes for each tumor state.

**Supplementary Table 5.** CollecTRI inferences for each tumor state.

**Supplementary Table 6.** Reactome pathway inferences for each tumor state. **Supplementary Table 7.** Moran’s I spatial autocorrelation results.

**Supplementary Table 8**. nAUC and statistical tests on scIDUC drug predictions.

**Supplementary Table 9.** Chemistry used for scRNA-seq for each sample.

